# Identification of Mitochondrial-Lipid Metabolism-Related Diagnostic Targets in Osteoarthritis: Integrative Multi-omics Data and Experimental Validation

**DOI:** 10.64898/2026.09.18.752655

**Authors:** Miao Huang, Xue Tan, Ji-Bin Yang, Li-Dan Yang, Jin Yang

**Affiliations:** Department of Orthopedic Surgery, Affiliated Hospital of Zunyi Medical University, Zunyi 563000, Guizhou Province, China; Department of Ophthalmology, Guizhou Aerospace Hospital (Affiliated Aerospace Hospital of Zunyi Medical University), Zunyi 563003, Guizhou Province, China

**Author notes:** Corresponding Author: Jin Yang, Department of Orthopedic Surgery, Affiliated Hospital of Zunyi Medical University, No.149 Dalian Road, Zunyi 563000, People’s Republic of China. Author Contributions: Miao Huang: Conceptualization, Methodology, Software, Formal analysis, Investigation, Writing – Original Draft, Visualization. Xue Tan: Validation, Formal analysis, Data Curation. Ji-Bin Yang: Methodology, Software, Validation. Li-Dan Yang: Resources, Data Curation. Jin Yang: Conceptualization, Resources, Writing – Review & Editing, Supervision, Project administration, Funding acquisition. All authors have read and agreed to the published version of the manuscript.

**Keywords:** Osteoarthritis, Mitochondrial-lipid metabolism, Hub genes, Machine learning, Diagnostic model

## Abstract

**Objective:** Osteoarthritis (OA) is a degenerative joint disease associated with metabolic dysregulation. This study aims to identify mitochondrial-lipid metabolism-related hub genes and explore their mechanisms in OA.

**Methods:** Training (GSE55235) and validation (GSE55457 and GSE82107) cohorts were retrieved from the Gene Expression Omnibus (GEO) database. Candidate genes were derived by overlapping differentially expressed genes, mitochondrial-lipid metabolism-related genes, and weighted gene co-expression network analysis module genes. Machine learning algorithms were applied to screen hub genes. The diagnostic performance, functional pathways, immune infiltration, regulatory networks, drug interactions, and molecular docking of the hub genes were systematically evaluated. An interleukin-1β-stimulated chondrocyte OA model was assayed for cell viability, interleukin-6 (IL-6), apoptosis, and hub-gene expression via Cell Counting Kit-8, enzyme-linked immunosorbent assay, flow cytometry, and reverse transcription-polymerase chain reaction.

**Results:** Twenty-five candidate genes were screened, from which six hub genes (alpha-methylacyl-CoA racemase [AMACR], uncoupling protein 2 [UCP2], acyl-CoA thioesterase 7 [ACOT7], aldehyde dehydrogenase 3 family member A2 [ALDH3A2], armadillo repeat containing X-linked 2 [ARMCX2], and monoamine oxidase A [MAOA]) were further identified. The six hub genes exhibited robust diagnostic ability in the training cohort, correlated with immune-cell infiltration, and participated in lipid-metabolism pathways. In the OA model, chondrocyte viability was reduced, while IL-6 levels and apoptosis rate were elevated; five hub genes were markedly upregulated, whereas MAOA was significantly downregulated.

**Conclusion:** Mitochondrial-lipid metabolism-related hub genes may serve as potential diagnostic biomarkers for OA, providing new insights into OA pathogenesis and potential therapeutic targets.

## Introduction

Osteoarthritis (OA) is a chronic degenerative joint disease that affects the entire joint. It arises from an imbalance between joint tissue repair and degradation, leading to degeneration and loss of articular cartilage, subchondral bone sclerosis, osteophyte formation, subchondral bone cysts, joint capsule fibrosis, synovial hyperplasia, and synovitis^[1]^.Among these pathological changes, cartilage degradation is considered the most important feature. OA most commonly affects the knee, hip, and hand joints, and represents a major cause of pain and physical dysfunction in the elderly^[2]^.Currently, OA remains incurable, and the efficacy of existing therapeutic approaches is limited. Current treatment primarily focuses on alleviating pain and other symptoms, as well as improving joint function. For patients with advanced-stage disease whose symptoms persist despite conservative treatment, joint replacement surgery is a viable option; however, the lifespan of artificial joint prostheses is limited^[3]^. Therefore, in-depth exploration of key genes involved in the initiation and progression of OA, as well as the identification of early diagnostic biomarkers with high sensitivity and specificity, may facilitate early intervention in OA and secure a more favorable therapeutic window for patients.

Mitochondria act as the primary nexus for lipid metabolism, extending far beyond their canonical function as cellular powerhouses by integrating fatty acid oxidation, de novo lipid synthesis, organelle dynamics, and redox homeostasis.Fatty acids primarily serve as substrates for ATP production in various organs of the human body and animals.^[4]^.The β-oxidation process, which is the most important component of fatty acid oxidation, occurs predominantly in the mitochondria of multiple organs, and energy generation is closely associated with mitochondrial β-oxidation. Mitochondria play a critical role in lipid oxidation and synthesis, and physical contacts between mitochondria and other organelles maintain the context-dependent balance required for cellular lipid homeostasis. Dysregulation of lipid homeostasis can exert disruptive effects on mitochondrial morphology and function^[5]^. Recent studies have established that mitochondrial dysfunction is a key factor in the pathogenesis of OA, promoting chondrocyte apoptosis, oxidative stress, and extracellular matrix degradation.^[6]^.Within joint tissues, mitochondria participate in regulating the activity of enzymes that control matrix remodeling, a process that ensures the continuous renewal and regeneration of cartilage. By modulating cellular metabolism, mitochondria influence the synthesis of molecules involved in maintaining the extracellular matrix (ECM), which is essential for joint integrity because it ensures that cartilage retains its elastic and load-bearing properties^[7]^. However, systematic studies on the mechanisms by which mitochondrial dysfunction and lipid metabolism disorders contribute to OA remain lacking. Therefore, further exploration of the core targets linking mitochondria and lipid metabolism to OA is expected to provide new directions for targeted therapy of OA.

Current studies indicate that the interplay between mitochondria and lipid metabolism plays an important role in diseases such as osteoarthritis (OA); however, the underlying mechanisms remain unclear. Therefore, further investigation into the role of mitochondrial lipid metabolism in OA is warranted. This study aims to identify key mitochondrial lipid metabolism-related hub genes that play critical roles in OA through bioinformatics analysis and *in vitro* experiments, and to systematically evaluate their specific regulatory mechanisms in OA, thereby providing a theoretical basis for the future treatment of OA and the development of targeted therapeutic agents.

## Materials and Methods

### Data Acquisition

The training and validation cohorts were both derived from the Gene Expression Omnibus (GEO) database (https://www.ncbi.nlm.nih.gov/geo/). For the training cohort^[8]^, synovial tissue samples from 10 patients with osteoarthritis (OA, Case group) and 10 healthy controls (Control group) were selected and profiled using microarray technology on the GPL96 platform. This dataset was employed for subsequent bioinformatics analyses. For the GSE55457 validation cohort^[8]^, synovial membrane samples from 10 patients with OA (Case group) and 10 healthy controls (Control group) were selected and measured on the GPL96 microarray platform. The GSE82107 validation cohort^[9]^ consisted of synovial biopsy samples from 10 patients with OA (Case group) and 7 non-arthritic controls (Control group), all profiled using the GPL570 microarray platform. These validation cohorts were used to confirm the expression of the identified hub genes, perform receiver operating characteristic (ROC) diagnostic analyses, and conduct external validation of the diagnostic model.

Mitochondria-related genes were retrieved from the Human MitoCarta3 database (http://www.broadinstitute.org/mitocarta). Lipid metabolism-related genes were obtained from gene sets in the Molecular Signatures Database (MSigDB, https://www.gsea-msigdb.org/gsea/msigdb) that were associated with the keywords “FATTY_ACID_METABOLISM”, “LIPID_METABOLIC”, “BILE_ACID_METABOLISM”, “CHOLESTEROL_HOMEOSTASIS”, and “ADIPOGENESIS”. An intersection was taken between the lipid metabolism-related gene sets and the mitochondria-related gene set, thereby generating a collection of mitochondrial-lipid metabolism-related genes, which was used for subsequent candidate gene screening.

### Differential Expression Analysis

To preliminarily screen for key genes implicated in the pathogenesis of OA, differential expression analysis was conducted on the training cohort using the limma package. Differentially expressed genes (DEGs) were identified with the filtering criteria of p-value < 0.05 and absolute log2 fold change (|log2FC|) > 0.5. A volcano plot was generated with the ggplot2 package to display all DEGs, and the top 10 upregulated and top 10 downregulated DEGs were annotated. In addition, the expression patterns of these top 10 upregulated and top 10 downregulated DEGs between the Control and Case groups were analyzed, and a heatmap was produced using the ComplexHeatmap package.

### Weighted Gene Co-expression Network Analysis (WGCNA)

To identify functional gene modules closely associated with the OA phenotype, a WGCNA was performed on the training cohort. First, hierarchical clustering of all samples was performed using Euclidean distances between gene expression profiles to detect and remove outlier samples. Then, the optimal soft-thresholding power was selected based on the scale-free topology fit index (R²), with the criterion that R² first exceeded 0.85. Subsequently, the gene co-expression similarities were transformed into a weighted adjacency matrix, which was further converted into a topological overlap matrix (TOM). Based on the TOM matrix, hierarchical clustering and dynamic tree cutting were applied to initially define modules, and modules were subsequently merged using a merge cut height of 0.25 according to the correlation of module eigengenes (MEs). Finally, the module eigengenes were used as representatives of module expression, and Pearson correlation coefficients were calculated between each module and the OA phenotype (OA vs. no-NA) to identify the modules that were significantly associated with the OA phenotype.

### Candidate Gene Identification

To obtain candidate genes that were differentially expressed in OA, involved in mitochondrial-lipid metabolism, and associated with OA-related WGCNA modules, an overlap analysis was performed among DEGs from the training cohort, mitochondrial-lipid metabolism genes, and WGCNA module genes. A Venn diagram generated by the ggvenn R package visualizes the intersection, and the intersecting genes are designated as candidate genes for subsequent analyses and hub-gene screening.

### Protein-Protein Interaction (PPI) and Functional Enrichment of Candidate Genes

To investigate the PPI relationships among the candidate genes, a PPI network was constructed using the Retrieval of Interacting Genes/Proteins (STRING) database (https://string-db.org/) with a combined score ≥ 300. The degree of each node was calculated, and the top 20 genes with the highest degrees were selected. A PPI network comprising these 20 genes and their interactions was then visualized as a static plot using the ggraph package in R. To explore the potential biological functions and molecular pathways in which the candidate genes might be involved, Gene Ontology (GO) enrichment analysis, including biological process (BP), cellular component (CC), and molecular function (MF) terms, as well as Kyoto Encyclopedia of Genes and Genomes (KEGG) pathway enrichment analysis, were performed using the clusterProfiler package. Enrichment results were filtered by a p-value < 0.05, from which the five most significant GO terms and the top 10 KEGG pathways (ranked by gene number) were displayed via the ggplot2 package. Additionally, a Sankey diagram was constructed to visualize the correspondence between candidate genes and enriched KEGG pathways.

### Hub Gene Selection Using Machine Learning Algorithms

To screen for hub genes among the candidate genes, three machine learning algorithms were applied. The Least Absolute Shrinkage and Selection Operator (LASSO) regression was performed using the glmnet package, with the optimal penalization parameter (λ) selected based on the minimum partial likelihood deviance; features with non-zero coefficients at λ were retained as contributors to OA classification. The Boruta algorithm was implemented with the Boruta package, which compared the importance between genuine features and shadow features based on random forest; a feature was considered important if its importance exceeded the maximum importance of the shadow attributes. The Support Vector Machine Recursive Feature Elimination (SVM-RFE) algorithm was conducted using the caret package for feature selection. During the process, features were ranked according to the SVM weight vector at each iteration, and the feature with the smallest ranking criterion was recursively removed. The subset that yielded the highest cross-validated accuracy was selected as the optimal feature set. Subsequently, a Venn diagram was generated using the ggvenn package to identify the overlapping genes among the three feature sets, and these overlapping genes were designated as hub genes for subsequent bioinformatics analyses.

### Evaluation and Diagnosis of Hub Genes

To evaluate the expression differences of the identified hub genes between the OA and Control groups, the Wilcoxon rank-sum test was performed separately in the training and two validation cohorts. The expression levels of each hub gene were compared between the OA and control samples in each dataset, with a significance threshold of p-value < 0.05. Subsequently, ROC curve analysis was conducted using the pROC package to assess the diagnostic discrimination ability of each hub gene. For each gene, the area under the curve (AUC) was computed to comprehensively evaluate diagnostic performance, with higher AUC values reflecting greater discriminatory ability.

### Diagnostic Model Construction and Evaluation

To develop a robust diagnostic model for OA based on the identified hub genes, eight commonly used machine learning algorithms were applied in the training cohort, including Generalized Linear Model (GLM), Naive Bayes (NB), Neural Network (NNET), LASSO regression, Random Forest (RF), SVM, K-Nearest Neighbors (KNN), and Elastic Net (ENET). OA status (OA vs. no-OA) was used as the response variable, and the expression levels of the hub genes were employed as explanatory variables. All models were constructed using the caret package in R. To interpret the model behavior and evaluate the contribution of each hub gene, the DALEX package was utilized to generate diagnostic plots, including cumulative residual distribution plots, boxplot-style residual distributions, and variable importance rankings. Multiple key metrics, including accuracy, F1-score, recall, specificity, precision, and sensitivity, were calculated for each model in the training cohort. The predictive performance of each model was further assessed based on two independent validation cohorts using the ROC curve analysis.

### Gene Set Enrichment Analysis (GSEA)

To explore the potential downstream signaling pathways associated with each hub gene, single-gene GSEA was performed in the training cohort using the clusterProfiler package. GSEA was performed for each hub gene using ranked gene lists based on Pearson correlation coefficients with all other genes in the genome-wide expression matrix, with the KEGG pathway gene sets obtained from MSigDB as the reference. Pathways with a p-value < 0.05 were considered significantly enriched and were visualized using the ggplot2 package.

### Immune Infiltration Analysis

To capture early immune abnormalities in OA and to quantify subtle alterations in multiple immune cell populations, single-sample Gene Set Enrichment Analysis (ssGSEA) was performed on the training cohort using the GSVA package. The enrichment scores of 28 immune cell phenotypes were calculated using the ssGSEA algorithm, and differences in immune infiltration between the OA and Control groups were evaluated via the Wilcoxon rank-sum test, with p-value < 0.05 considered statistically significant. The results were visualized using the ggplot2 package to display the score distribution across groups. Furthermore, to explore the associations between the identified hub genes and immune cell infiltration, Pearson correlation analysis was conducted between the expression levels of the hub genes and the enrichment scores of each immune cell type.

### Chromosomal and Subcellular Localization

To examine the chromosomal distribution of the identified hub genes, their genomic locations were mapped onto human chromosomes using the RCircos package. A circular ideogram was generated to visualize the chromosomal positions of each hub gene. To further investigate the subcellular localization of the proteins encoded by the hub genes, the gene sequences were submitted to the mRNALocater database (http://bio-bigdata.cn/mRNALocater) for subcellular localization prediction. Additionally, to explore the functional associations among the hub genes and their related genes, a functional association network was constructed using the GeneMANIA database (http://genemania.org). This network integrated multiple association types, thus elucidating the functional landscape of the hub genes and their associated partners.

### Regulatory Network Construction

To investigate the upstream regulators of the hub genes at the transcriptional and post-transcriptional levels, the miRNet platform (https://www.mirnet.ca) was employed to identify potential microRNAs (miRNAs) and transcription factors (TFs) targeting the hub genes. Specifically, TF predictions were obtained from the ChIP-X Enrichment Analysis (ChEA) database, while miRNA-hub gene interaction pairs were retrieved using the miRNA target prediction module. All predicted regulatory relationships were subsequently loaded into Cytoscape software for network visualization.

### Drug Prediction and Molecular Docking

To identify potential candidate drugs targeting the hub genes, drug prediction analysis was performed using the Drug Signature Database (DSigDB; https://dsigdb.tanlab.org). After standardizing drug names and filtering for candidate drugs with more than 3 target genes, the drug– gene interaction network was constructed. To determine the binding affinities between candidate drugs and their target proteins, up to three drugs were selected for each hub gene and molecular docking was conducted via the CB-Dock2 platform (https://cadd.labshare.cn/cb-dock2). The corresponding chemical structures and physicochemical properties of the candidate drugs were retrieved from the PubChem database (https://pubchem.ncbi.nlm.nih.gov) and shown in supplementary Table 1. The sequence information of the hub genes was retrieved from the UniProt database (https://www.uniprot.org), while their three-dimensional structural models were obtained from AlphaFold (https://alphafold.ebi.ac.uk).

### Cell Culture and OA Model Establishment

Human primary chondrocytes (iCell Bioscience Inc, Shanghai, China) were cultured in Dulbecco’s Modified Eagle’s Medium (DMEM)/Nutrient Mixture F-12 medium^[10]^ (iCell Bioscience Inc) supplemented with 10% fetal bovine serum (FBS, Servicebio Technology Co., Ltd, Wuhan, China) and 1% penicillin/streptomycin (Servicebio Technology Co., Ltd) at 37°C in an atmosphere containing 5% CO₂. Chondrocytes were divided into Control and OA groups, with the OA group stimulated with 10 ng/mL interleukin-1β (IL-1β, Servicebio Technology Co., Ltd) for 24 hours to establish an *in vitro* OA model^[11]^.

### Cell Viability Assay

Cell viability was evaluated using the Cell Counting Kit 8 (CCK-8) assay. A 100 μL chondrocyte suspension was seeded into 96-well plates at a density of 1×10³ cells per well and cultured to 80% confluence. After 24 h of interleukin-1β (IL-1β) treatment, the culture medium was gently aspirated, and the cells were washed twice with Dulbecco’s phosphate-buffered saline (DPBS). Then, 10 μL of CCK-8 solution (Beyotime Biotechnology Co., Ltd., Shanghai, China) was added to each well, and the plates were incubated for 2 h. The absorbance at 450 nm was measured using a microplate reader.

### Enzyme-Linked Immunosorbent Assay (ELISA)

The concentration of interleukin-6 (IL-6) in cell supernatants was measured using a Human IL-6 ELISA Kit (Beyotime Biotechnology Co., Ltd.) according to the manufacturer’s instructions. Briefly, samples and standards were added to the pre-coated plates, followed by incubation with Horseradish peroxidase (HRP)-conjugated detection antibody. After washing, substrate solutions were added, and the optical density was measured at 450 nm.

### Apoptosis Detection by Flow Cytometry

Chondrocyte apoptosis was assessed using an Annexin V-FITC Apoptosis Detection Kit (Beyotime Biotechnology Co., Ltd.). Briefly, chondrocytes were harvested, washed with phosphate-buffered saline (PBS), and resuspended in 195 μL of Annexin V-Fluorescein Isothiocyanate (FITC) binding buffer. Subsequently, 5 μL of Annexin V-FITC and 10 μL of propidium iodide (PI) were added, followed by incubation at room temperature in the dark for 10– 20 minutes. Apoptotic cells were immediately analyzed using a flow cytometer.

### Reverse Transcription Quantitative Polymerase Chain Reaction (RT-qPCR)

Total RNA was extracted from cells using TRIzol reagent (Servicebio Technology Co., Ltd.), and cDNA was synthesized using the Hiscript II QRT Supermix for qPCR (Vazyme Biotech Co., Ltd, Nanjing, China) according to the manufacturer’s protocol. The qPCR was performed using 2 × ChamQ Universal SYBR qPCR Master Mix (Vazyme Biotech Co., Ltd) in 20 μL reaction system. Glyceraldehyde-3-phosphate dehydrogenase (GAPDH) was employed as the internal reference gene, and the corresponding primer sequences were provided in Supplementary Table 2. Relative gene expression levels were calculated using the 2^⁻ΔΔCt^ method.

### Statistical Analysis

All statistical analyses were performed using R software and GraphPad Prism (version 10.0). For the bioinformatics analyses, expression differences were evaluated using the Wilcoxon rank-sum test. Correlation analyses were performed using Spearman’s rank correlation. All tests were two-sided, and a p-value < 0.05 was considered statistically significant unless otherwise specified. For the *in vitro* experiments, all data were presented as mean ± standard deviation (SD). Comparisons between two groups were performed using a t-test, and a p-value < 0.05 was regarded as statistically significant.

## Results

### Candidate Genes Identification

A total of 422 mitochondrial-lipid metabolism-related genes were obtained by intersecting the lipid metabolism-related gene set with the mitochondrial-related gene set (Figure 1A). In addition, 2,584 DEGs were identified in the training cohort, of which 1,259 were upregulated and 1,325 were downregulated. The overall distribution of these DEGs was visualized using a volcano plot (Figure 1B), and a heatmap displayed the distinct expression patterns of the top 10 upregulated and top 10 downregulated DEGs across the Case and Control groups (Figure 1C).

**Figure 1.**
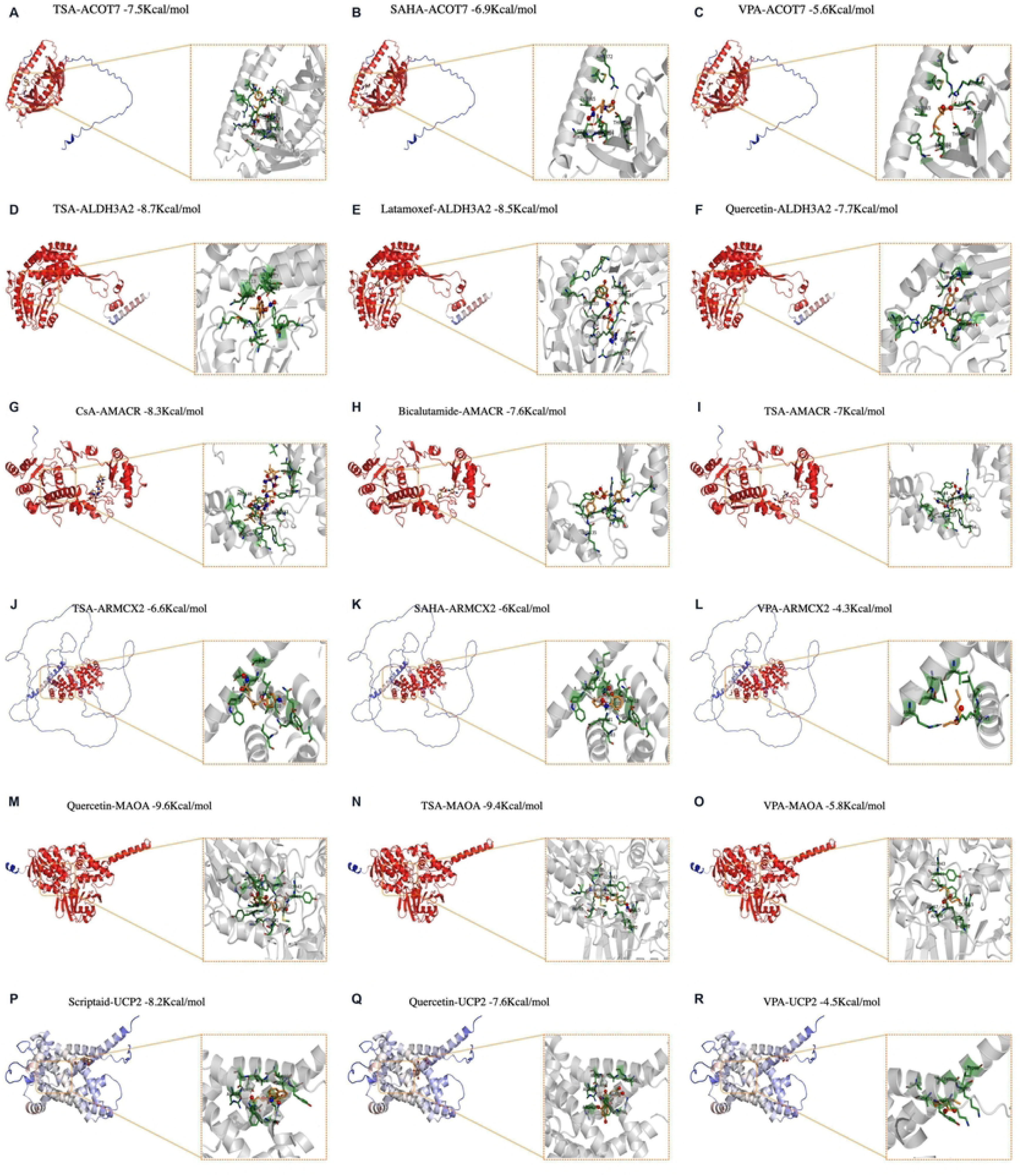
Screening of candidate genes via integrative bioinformatics analysis. (A) Mitochondrial lipid metabolism-related gene identification. The blue circle represents lipid-metabolism-related genes, and the orange circle represents mitochondrial-related genes. The overlapping area indicates mitochondrial-lipid metabolism-related genes. (B) Volcano plot of differentially expressed genes (DEGs) between the osteoarthritis (OA, Case group) and Control group. Red dots denote significantly upregulated genes, blue dots represent significantly downregulated genes, and grey dots indicate genes with no significant expression change. Dashed lines mark the screening threshold of DEGs. (C) Density distribution and expression heatmap of the top 10 upregulated and top 10 downregulated DEGs. The upper panel displays the density distribution of gene expression. The lower heatmap shows normalized expression levels of DEGs across samples. Green bars correspond to the Control group, and red bars correspond to the Case group. The color gradient reflects normalized gene expression values, where red indicates high expression, and green indicates low expression. (D) Sample dendrogram and trait heatmap for sample clustering. The y-axis represents clustering height. The bottom-row trait heatmap illustrates sample group information; varying red shades distinguish different sample groups. (E) Determination of soft-thresholding power for weighted gene co-expression network construction. The left panel shows scale-independence analysis; the x-axis corresponds to soft-thresholding, and the y-axis represents scale-free topology model fit (signed R²). The red horizontal line indicates the threshold of R² = 0.85. The right panel exhibits mean connectivity under different soft-thresholding powers. The x-axis is soft-thresholding, and the y-axis denotes mean connectivity. (F) Gene cluster dendrogram for module identification. The y-axis indicates clustering height. The bottom color bar assigns each gene to its corresponding co-expression module; different colors represent distinct gene modules. (G) Heatmap of module-trait relationships between gene modules and clinical phenotypes. Each cell contains the correlation coefficient and corresponding p-value in parentheses. The color gradient reflects correlation strength: red denotes positive correlation, and blue denotes negative correlation. (H) Candidate gene identification. The blue circle represents DEGs, the green circle represents mitochondrial-lipid metabolism-related genes, and the pink circle represents WGCNA module genes. Genes in the overlapping central region are defined as candidate genes.

In WGCNA, hierarchical clustering of all samples revealed a homogeneous distribution without obvious outliers, and a tendency toward grouping between the OA and control samples was observed (Figure 1D). The optimal soft-thresholding power was determined as β = 7, at which point the R² first exceeded 0.85 and the mean connectivity reached a moderate level (Figure 1E). A total of 17 distinct modules were identified via dynamic tree cutting and module merging (Figure 1F). Correlation analysis between module eigengenes and the OA phenotype showed that the MEyellow module was most significantly positively correlated with OA (r = 0.950, p-value < 0.05), whereas the MEbrown module exhibited a significant negative correlation (r = -0.932, p-value < 0.05) (Figure 1G). These two modules were therefore considered key modules, containing a total of 1,867 module genes.

Intersection analysis was performed among the DEGs in the training cohort, mitochondrial lipid metabolism-related genes, and module genes derived from WGCNA, yielding a total of 25 overlapping genes, which were designated as final candidate genes for subsequent hub gene screening (Figure 1H).

### PPI Network and Functional Enrichment of Candidate Genes

A PPI network was built using the STRING database to investigate the protein-level relationships of the candidate genes (Figure 2A). Based on node degree, the top 20 genes were further selected to generate a network, which consisted of 20 nodes and 38 edges (Figure 2B). The top 10 interacting pairs with the highest combined score were mitochondrial ribosomal protein S34 (MRPS34)–mitochondrial ribosomal protein S33 (MRPS33) (0.992), peptidylprolyl isomerase F (PPIF)–cytochrome c, somatic (CYCS) (0.983), acyl-CoA dehydrogenase long chain (ACADL)– carnitine palmitoyltransferase 2 (CPT2) (0.982), monoamine oxidase A (MAOA)–aldehyde dehydrogenase 3 family member A2 (ALDH3A2) (0.957), ALDH3A2–aldo-keto reductase family 1 member B10 (AKR1B10) (0.951), ALDH3A2–propionyl-CoA carboxylase subunit alpha (PCCA) (0.850), aldo-keto reductase family 7 member A2 (AKR7A2)–AKR1B10 (0.781), ACADL–PCCA (0.664), CYCS–uncoupling protein 2 (UCP2) (0.619), and PCCA–glutaryl-CoA dehydrogenase (GCDH) (0.564).

**Figure 2.**
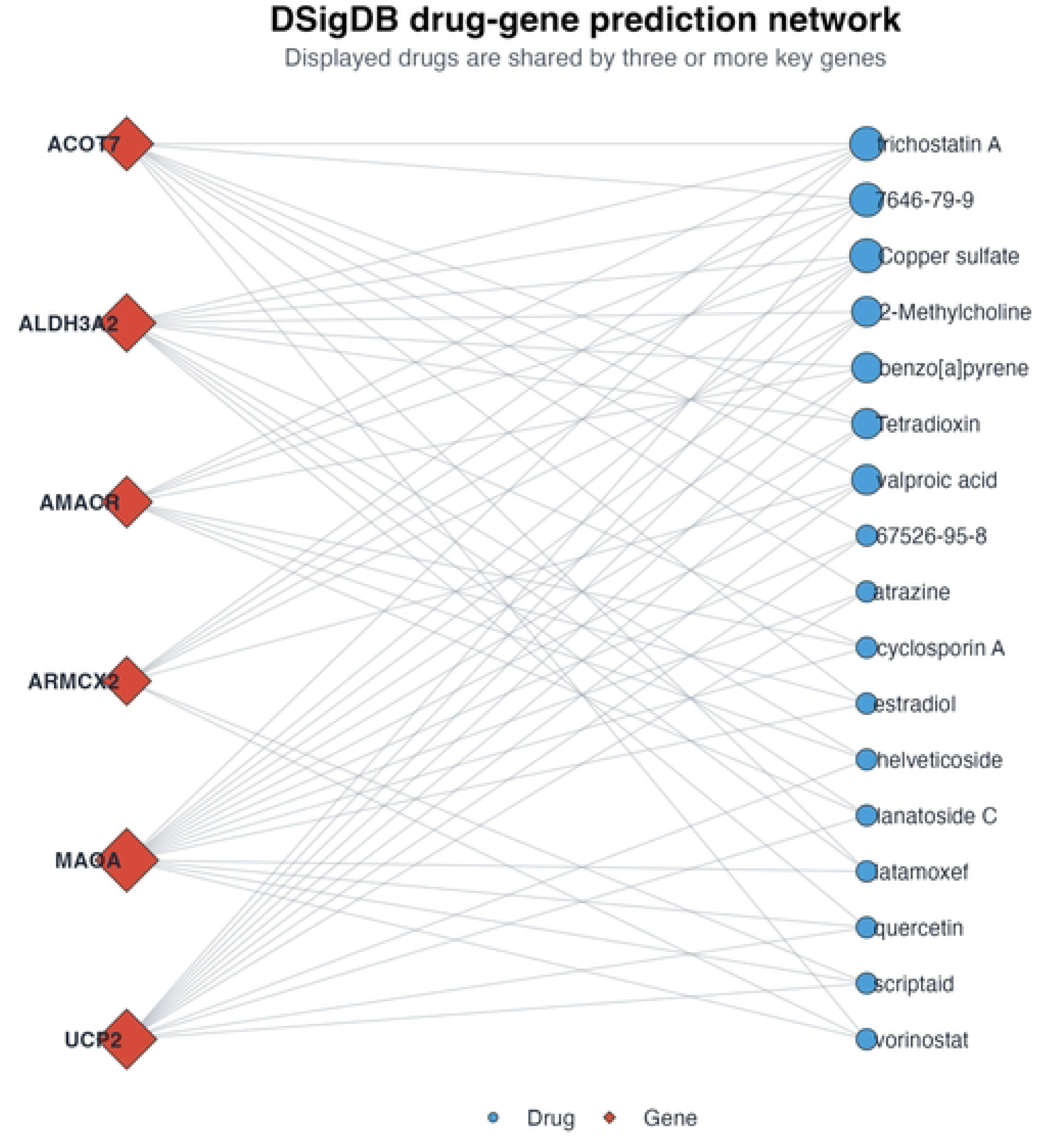
Protein-protein interaction (PPI) network and functional enrichment analysis of candidate genes. (A) Original PPI network output from the Retrieval of Interacting Genes/Proteins (STRING) database. Nodes denote proteins, and line thickness reflects the confidence level of each PPI. Different node colors represent distinct proteins. (B) PPI network of top 20 candidate genes ranked by degree value. Each circle represents one protein node, and node size and color gradient correspond to node degree. Lines between nodes stand for PPIs. (C) Top five Gene Ontology (GO) enrichment terms of candidate genes. Horizontal bar chart displays the top five enriched GO terms for biological process (BP, red), cellular component (CC, teal), and molecular function (MF, beige). The x-axis represents the number of enriched genes, and the y-axis lists corresponding GO terms. (D) Kyoto Encyclopedia of Genes and Genomes (KEGG) pathway enrichment analysis of candidate genes. The x-axis indicates the number of enriched genes, and the y-axis represents pathway names. Dot size corresponds to gene count, and dot color reflects the p-value. (E) Sankey diagram of the top 10 KEGG pathways. The left Sankey diagram displays the mapping relationships between candidate genes and the top 10 enriched KEGG pathways. The right bubble plot presents KEGG enrichment statistics; the x-axis represents the rich factor; dot color corresponds to the adjusted p-value; dot size indicates the number of enriched genes within each pathway.

GO enrichment analysis was conducted for the 25 candidate genes. The top five enriched terms in BP mainly involved small molecule catabolic process, fatty acid metabolic process, organic acid catabolic process, carboxylic acid catabolic process, and cellular catabolic process. For CC, the significantly enriched terms included peroxisome, microbody, mitochondrial protein-containing complex, organellar small ribosomal subunit, and mitochondrial small ribosomal subunit. In terms of MF, the dominant enriched terms were isomerase activity, amide binding, sulfur compound binding, fatty-acyl-CoA binding, and fatty acid derivative binding (Figure 2C). The top 10 enriched KEGG pathways were fatty acid degradation, tryptophan metabolism, histidine metabolism, tyrosine metabolism, valine, leucine and isoleucine degradation, arginine and proline metabolism, fatty acid metabolism, lysine degradation, glycerolipid metabolism, and the peroxisome proliferator-activated receptor (PPAR) signaling pathway (Figure 2D-E).

### Identification and Validation of Hub Genes

Three machine-learning algorithms were applied to screen feature genes. For LASSO regression, the optimal λ value was determined as 0.0041 according to the minimal partial likelihood deviance, and eight feature genes with non-zero regression coefficients were retained (Figure 3A-B). The Boruta algorithm identified 17 confirmed feature genes based on variable importance (Figure 3C), and SVM-RFE obtained 12 feature genes at the highest classification accuracy (Figure 3D). The Venn diagram was used to acquire the intersection of feature genes screened by the three algorithms, and six hub genes were finally obtained for subsequent analysis (Figure 3E). The six hub genes were alpha-methylacyl-CoA racemase (AMACR), UCP2, acyl-CoA thioesterase 7 (ACOT7), ALDH3A2, armadillo repeat containing X-linked 2 (ARMCX2), and MAOA. The expression levels and ROC curves of the six hub genes were compared between the OA and control samples in the training cohort and two external validation cohorts. In the training cohort, AMACR, UCP2, ACOT7, ALDH3A2, and ARMCX2 were significantly upregulated, whereas MAOA showed significantly decreased expression in OA samples (Figure 3F). In the GSE55457 validation cohort, AMACR, UCP2, ACOT7, ALDH3A2, and ARMCX2 maintained significant upregulation in OA samples(Figure 3G). In the GSE82107 validation cohort, only ACOT7 and ARMCX2 were significantly upregulated in OA samples (Figure 3H).

**Figure 3.**
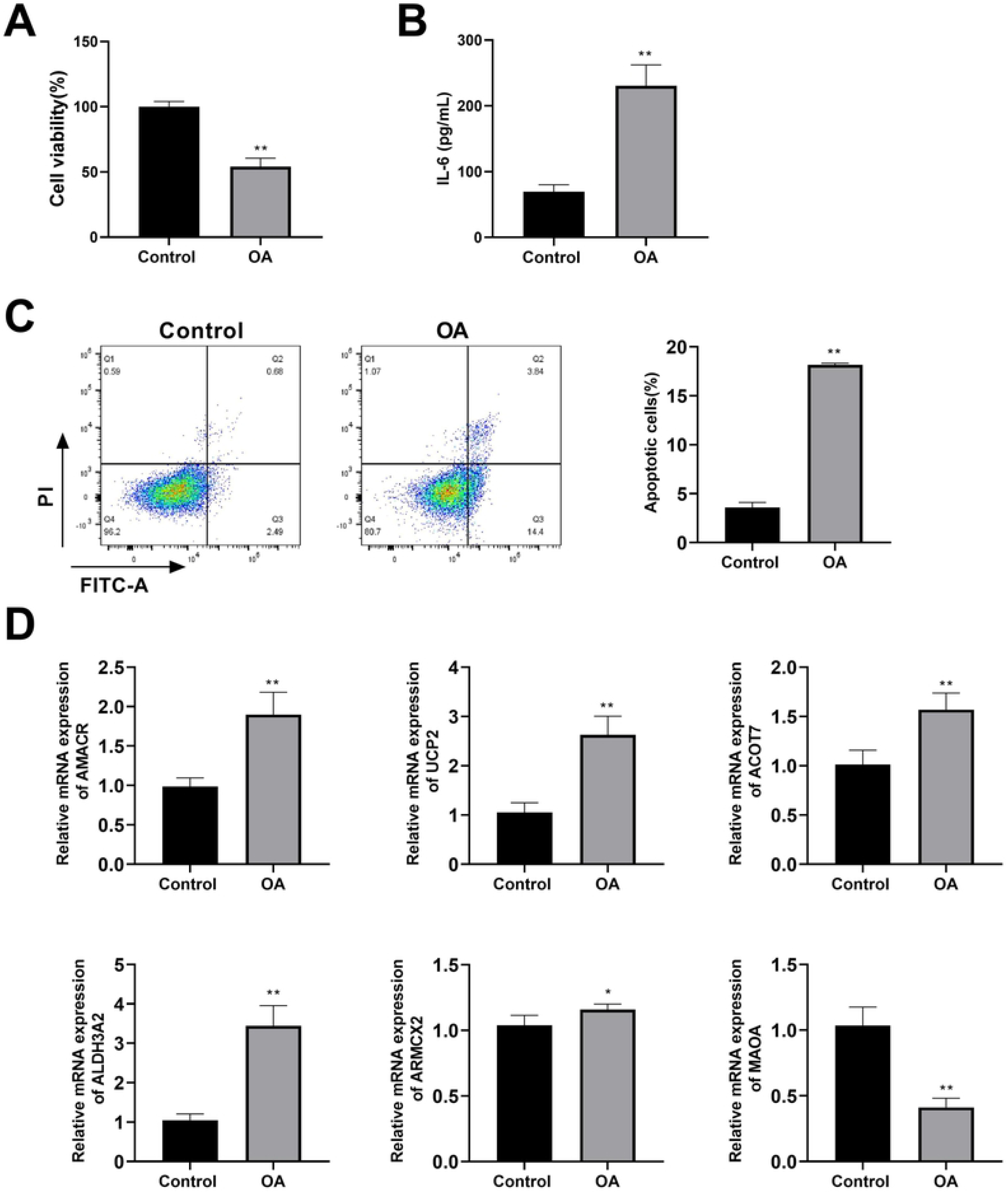
Identification and validation of hub genes. (A) Cross-validation curve for Least Absolute Shrinkage and Selection Operator (LASSO) regression. The x-axis shows logλ, while the y-axis indicates partial likelihood deviance derived from cross-validation. Colored lines display the model deviance corresponding to different numbers of retained variables. The vertical dashed line corresponds to λ.min. The legend at the top labels the number of variables matched to each color curve. (B) Coefficient profile plot of LASSO regression. Each coloured line displays the changing coefficient of an individual gene along with varying log λ. The vertical dashed line indicates the optimal λ value. (C) Variable importance distribution of feature genes calculated by the Boruta algorithm. Yellow boxes represent confirmed genes; shadowMax, shadowMean, and shadowMin denote shadow feature genes. The x-axis lists gene names, and the y-axis shows gene importance. (D) Accuracy curve of Support Vector Machine Recursive Feature Elimination (SVM-RFE) recursive feature elimination. The vertical dashed line indicates feature genes with the maximum classification accuracy. The x-axis is the number of input genes, and the y-axis represents classification accuracy. (E) Identification of hub genes. The pink circle denotes LASSO-feature genes, the purple circle illustrates SVM-RFE-feature genes, and the green circle shows Boruta-feature genes. (F) Expression comparison of hub genes in the training cohort. (G) Expression comparison of hub genes in the GSE82107 validation cohort. (H) Expression comparison of hub genes in the GSE55457 validation cohort. Box-plot analysis of hub-gene expression between OA (red) and control (blue) samples. The x-axis lists hub genes, and the y-axis represents gene expression values. Asterisks indicate statistical significance between the two groups;* p < 0.05; ** p < 0.01; *** p < 0.001; **** p < 0.0001; ns indicates no significant difference. (I) Receiver operating characteristic (ROC) analyses of hub genes in the training cohort. (J) ROC analyses of hub genes in the GSE55457 validation cohort. (K) ROC analyses of hub genes in the GSE82107 validation cohort. The x-axis represents 1-Specificity, and the y-axis represents Sensitivity. Each coloured line corresponds to one hub gene.

ROC curve analyses were conducted to assess the diagnostic performance of the six hub genes in distinguishing OA samples from controls. In the training cohort, all six genes exhibited strong discriminatory ability, with AUC values exceeding 0.90 (Figure 3I). In the GSE55457 validation cohort, AMACR, UCP2, ACOT7, ALDH3A2, and ARMCX2 maintained favorable diagnostic efficiency (AUC > 0.80), whereas MAOA showed a markedly lower AUC of 0.540 (Figure 3J). In the GSE82107 validation cohort, ACOT7, ARMCX2, and UCP2 yielded acceptable AUC values (AUC > 0.60), whereas AMACR, MAOA, and ALDH3A2 demonstrated limited diagnostic potential, with AUCs below 0.60 (Figure 3K). The AUC values, 95% confidence intervals (CIs), and accuracy of the six hub genes in the training and validation cohorts were presented in Supplementary Table 3.

### Assessment of Eight Machine-Learning Classifiers Derived from Six Hub Genes for OA Diagnosis

Eight machine-learning models were constructed based on six hub genes for OA diagnosis. Reverse cumulative distribution curves of absolute residuals were used to assess the prediction deviation of eight machine-learning models. The GLM and NB curves dropped sharply at low absolute residual values, indicating that most samples yielded small prediction errors (Figure 4A). Boxplots of absolute residuals revealed that the interquartile boxes for GLM and NB were barely visible, reflecting minimal prediction variability among most samples (Figure 4B). Permutation-based feature importance analysis was performed to evaluate the contribution of six hub genes to prediction performance across eight machine-learning models. Substantial differences in feature-importance ranking were observed among different algorithms. MAOA exhibited high predictive importance across most models. By contrast, ARMCX2 showed relatively low importance across most models, indicating a limited contribution to model prediction (Figure 4C). ROC curves revealed that RF, SVM, LASSO, ENET, KNN, NNET, and NB achieved an AUC value of 1.00, and the GLM model yielded an AUC of 0.98. These results suggested excellent discriminatory ability of the constructed models in the training cohort (Figure 4D). Most models achieved near-optimal performance across these evaluation indicators in the training cohort (Figure 4E). The predictive performance of eight machine-learning models was further assessed in two independent validation datasets. In the GSE55457 validation cohort, SVM (AUC = 1.00), ENET (AUC = 0.92), LASSO (AUC = 0.87), NNET (AUC = 0.74), NB (AUC = 0.70) and RF (AUC = 0.605) attained moderate-to-good diagnostic capacity (Figure 4F, Figure 4H). In the GSE82107 validation cohort, GLM (AUC = 0.85), ENET (AUC = 0.771), NB (AUC = 0.771), LASSO (AUC = 0.714), NNET (AUC = 0.700), KNN (AUC = 0.686) and SVM (AUC = 0.643) achieved moderate-or-higher diagnostic performance (Figure 4G-H).

**Figure 4.**
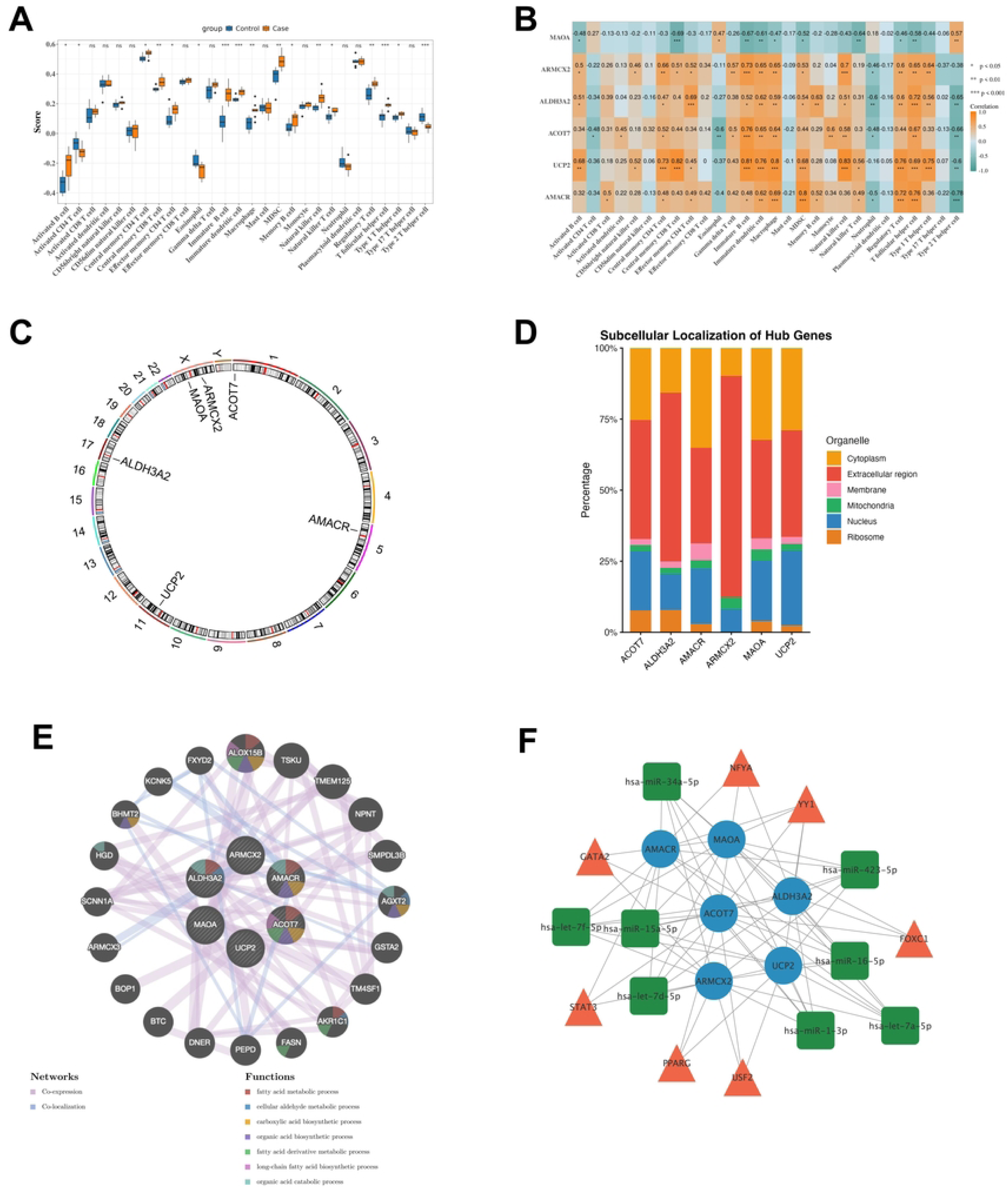
Assessment of eight machine-learning classifiers derived from hub genes for OA diagnosis. (A) Reverse cumulative distribution curves of absolute residuals for eight machine-learning models. The x-axis represents the absolute residual magnitude, and the y-axis shows the sample proportion. Curves dropping rapidly on the left represent models with small prediction errors for most samples. (B) Boxplots of absolute residuals across eight machine-learning models. The x-axis represents absolute residual values. Red dots indicate the root-mean-square error of residuals (RMSE). (C) Permutation-based feature importance of six hub genes in eight machine-learning models. The x-axis shows the increase in RMSE after feature permutation; larger values indicate greater predictive importance of the corresponding gene. Error bars denote the variability from repeated permutation. Each row corresponds to one machine-learning model. (D) ROC curves of eight machine-learning models in the training cohort. The x-axis represents the false-positive rate (1-specificity), and the y-axis shows the true-positive rate (sensitivity). (E) Performance evaluation of eight machine-learning models in the training cohort. The left-upper panel shows ROC curves for eight machine-learning algorithms. The right-upper panel presents the comparison of AUC values among different models. The bottom panel displays multiple evaluation metrics, including accuracy, F1-score, recall, specificity, AUC, precision, and sensitivity. The x-axis represents different models, and the y-axis corresponds to score values. (F) ROC curves of eight machine-learning models in the GSE55457 validation cohort. (G) ROC curves of eight machine-learning models in the GSE82107 validation cohort. The x-axis corresponds to false-positive rate (1-specificity), and the y-axis corresponds to true-positive rate (sensitivity). (H) AUC trajectory of eight machine-learning models across the training and two validation cohorts. Line plot illustrates the AUC variation of each machine-learning model among the training cohort, validation cohort 1 (GSE55457), and validation cohort 2 (GSE82107). The x-axis indicates different cohorts, and the y-axis represents AUC values. Each colored line stands for one machine-learning algorithm.

### Functional Annotation of Hub Genes by GSEA

GSEA was performed to explore the potential biological pathways associated with hub genes. For AMACR, the top six enriched pathways included asthma, type I diabetes mellitus, allograft rejection, lysosome, cell adhesion molecules (CAMs), and spliceosome (Figure 5A). For UCP2, the six most enriched pathways included antigen processing and presentation, viral myocarditis, CAMs, ribosome, spliceosome, and lysosome (Figure 5B). For ACOT7, the top six enriched pathways consisted of viral myocarditis, CAMs, ribosome, oxidative phosphorylation, spliceosome, and lysosome (Figure 5C). For ALDH3A2, the six most significantly enriched pathways were Fc gamma R-mediated phagocytosis, spliceosome, leishmania infection, Parkinson disease, oxidative phosphorylation, and lysosome (Figure 5D). For ARMCX2, the top six enriched pathways comprised CAMs, spliceosome, Parkinson disease, other glycan degradation, oxidative phosphorylation, and lysosome (Figure 5E). For MAOA, the top six enriched pathways were asthma, graft-versus-host disease, antigen processing and presentation, CAMs, spliceosome, and ribosome (Figure 5F). The normalized enrichment score (NES), adjusted p-value (padj), set size, and enrichment direction of the top 10 enriched pathways for each hub gene were summarized in Supplementary Table 4.

**Figure 5.**
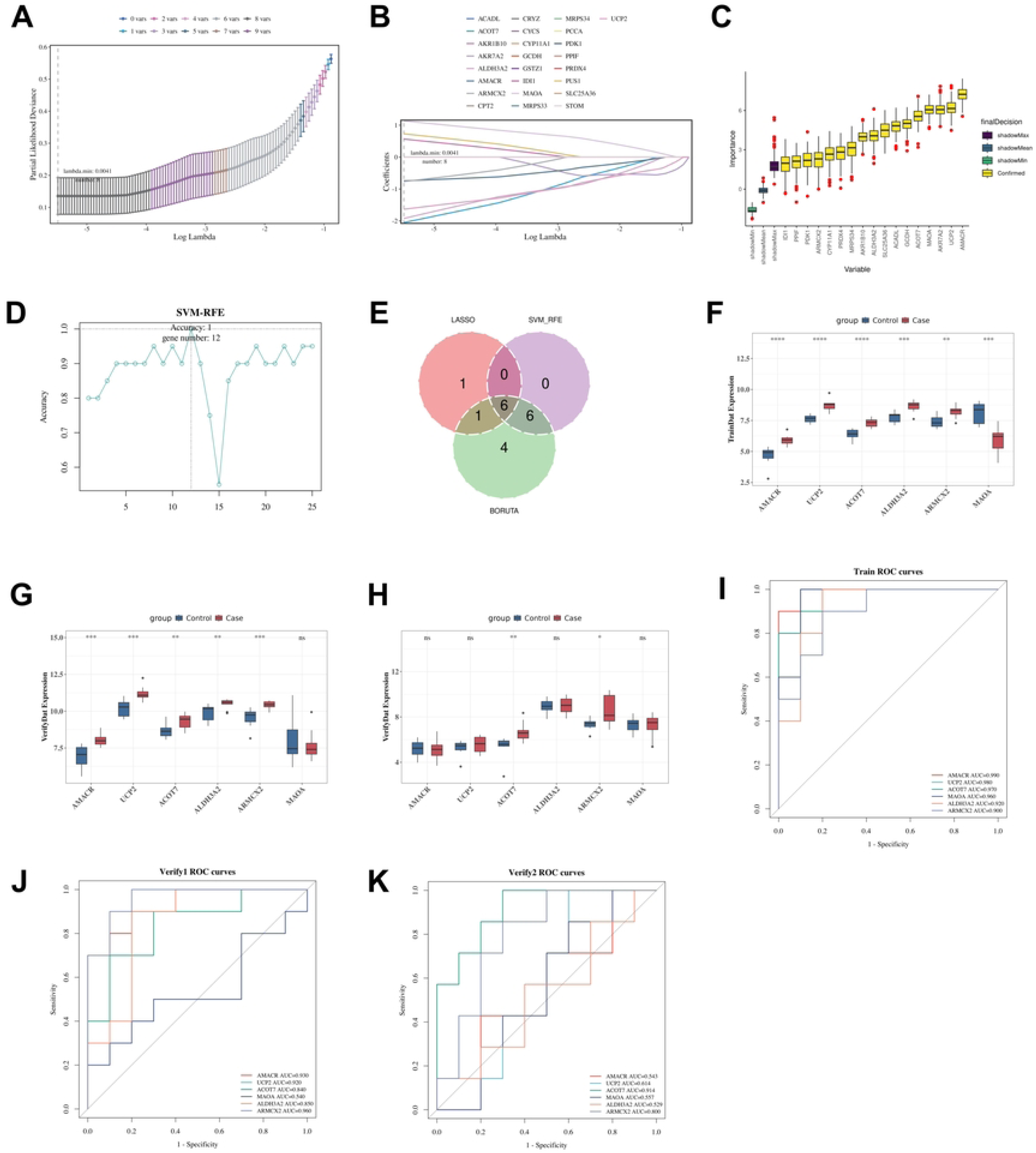
Gene set enrichment analysis (GSEA) of hub genes. (A) Alpha-methylacyl-CoA racemase (AMACR). (B) Uncoupling protein 2 (UCP2). (C) Acyl-CoA thioesterase 7 (ACOT7). (D) Aldehyde dehydrogenase 3 family member A2 (ALDH3A2). (E) Armadillo repeat containing X-linked 2 (ARMCX2). (F) Monoamine oxidase A (MAOA). The colored curves represent distinct functional pathways, and the vertical axis indicates the enrichment score. The vertical lines mark the genes enriched in the respective pathways. In the gray area below, the horizontal axis corresponds to genes ranked by their correlation, and the vertical axis shows the corresponding ranked list metric.

### Comprehensive Analysis of Hub Genes

Immune cell infiltration patterns were compared between OA and control samples. Significant differences in immune-cell infiltration scores were detected for multiple immune cell subsets between the two groups. The infiltration scores of activated B cells, central memory CD4 T cells, central memory CD8 T cells, effector memory CD4 T cells, immature B cells, immature dendritic cells, macrophages, myeloid-derived suppressor cells (MDSCs), memory B cells, natural killer cells, natural killer T cells, regulatory T cells, T follicular helper cells, and Type 1 T helper cells were markedly higher infiltration, whereas activated CD4 T cells, eosinophil, and Type 2 T helper cells were markedly decreased infiltration in OA samples (Figure 6A). Analysis of the correlations between the six hub gene expression levels and the infiltration scores of 28 immune cell types revealed that all six genes were significantly associated with most immune cell types. A generally consistent correlation pattern was observed among AMACR, UCP2, ACOT7, ALDH3A2, and ARMCX2, whereas MAOA exhibited an opposite correlation pattern with the other five hub genes (Figure 6B).

**Figure 6.**
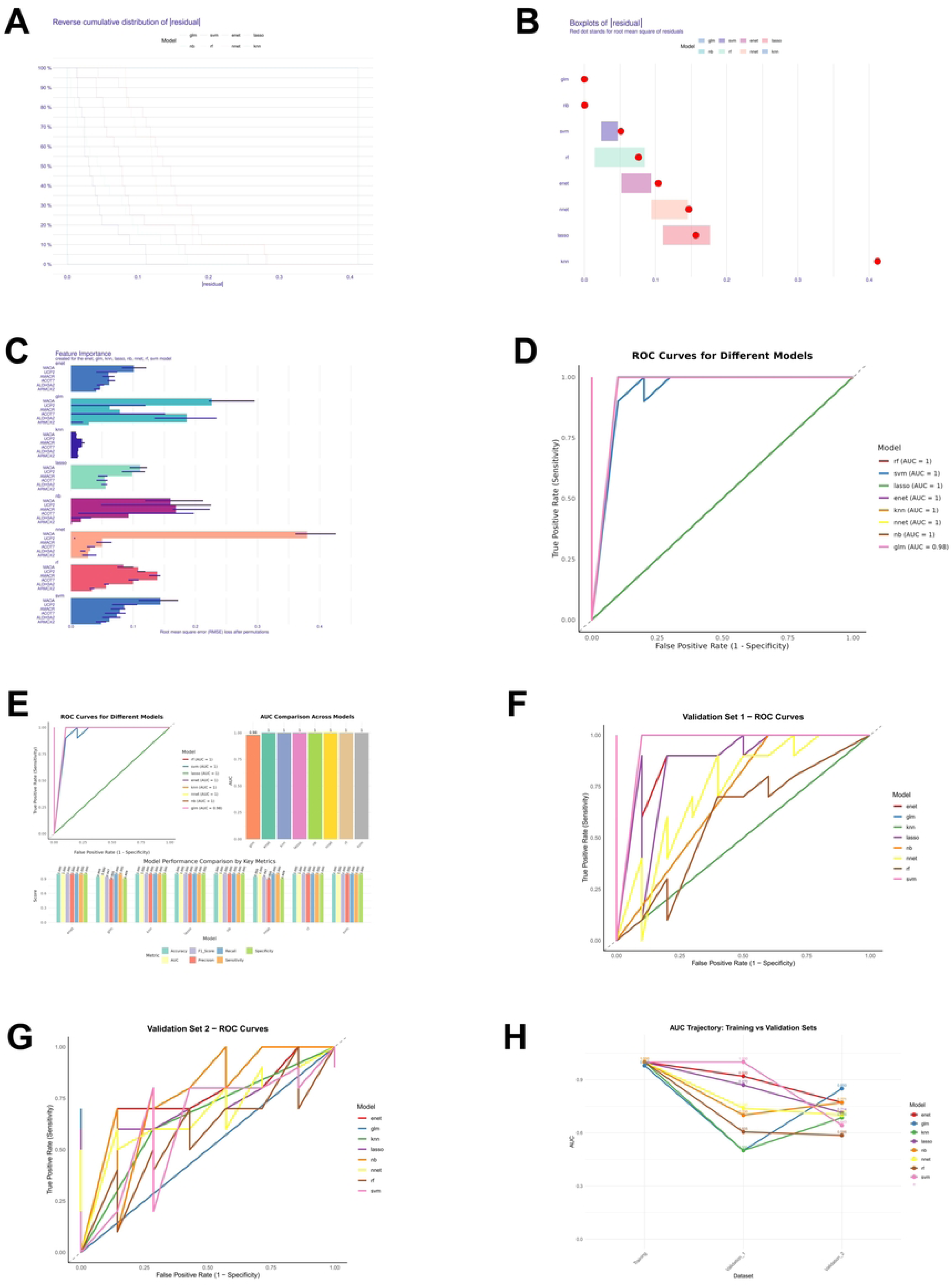
Integrated analysis of immune correlations, genomic localization, and regulatory networks of hub genes. (A) Differences in immune-cell infiltration scores between Control and Case groups. Blue represents the Control group, and orange represents the Case group. The x-axis shows the names of 28 types of immune cells, and the y-axis shows the score. “ns” indicates no significant difference. (B) Correlation between hub-gene expression and immune-cell infiltration. Rows stand for the hub genes, and columns correspond to distinct immune cell subtypes. The color gradient reflects the correlation coefficient value and direction: orange color indicates positive correlation, while green color represents negative correlation. Asterisks denote statistical significance: *p < 0.05, **p < 0.01; ***p < 0.001. (C) Chromosomal distribution of hub genes. Numbers on the outer ring represent human autosomes, while X and Y indicate sex chromosomes. Labels inside the circle correspond to the chromosomal location of each hub gene. (D) Predicted subcellular localization of hub genes-encoded proteins. The x-axis lists the six hub genes, and the y-axis shows the percentage of localization probability. (E) Functional association network of hub genes. The size of each node reflects its degree of connection, and the larger the node, the higher its correlation. (F) The microRNA (miRNA)-mRNA-transcription factor (TF) network. Blue circles represent hub gene mRNAs, green rectangles represent miRNAs, and orange triangles represent TFs.

The chromosomal localization analysis revealed that ACOT7, AMACR, UCP2, and ALDH3A2 were distributed on chromosomes 1, 5, 11, and 17, respectively, with MAOA and ARMCX2 both located on the X chromosome (Figure 6C). Regarding subcellular distribution, the proteins encoded by the hub genes were predominantly localized to the cytoplasm, extracellular region, nucleus, and mitochondria. Additionally, except for ARMCX2, the remaining five hub gene-encoded proteins were also detected in the cell membrane and ribosomes (Figure 6D).

A gene-gene interaction network revealed the potential interactions among candidate genes. Co-expression and co-localization constituted the main interaction modes within this network. Functional annotation suggested that these interacting genes were predominantly enriched in lipid-related biological processes (e.g., fatty acid metabolic process, cellular aldehyde metabolic process, and carboxylic acid biosynthetic process) (Figure 6E).

TF and miRNA regulators of hub genes were predicted and integrated to build a TF-mRNA-miRNA regulatory network. Transcription factors (e.g., nuclear transcription factor Y subunit alpha [NFYA], forkhead box C1 [FOXC1], and GATA binding protein 2 [GATA2]) and multiple miRNAs (e.g., hsa-miR-34a-5p, hsa-let-7f-5p, hsa-miR-15a-5p) were predicted to target the six hub genes. Widespread cross-regulatory links among network nodes suggested that hub genes were controlled by intricate TF-miRNA regulatory axes (Figure 7F).

**Figure 7.**
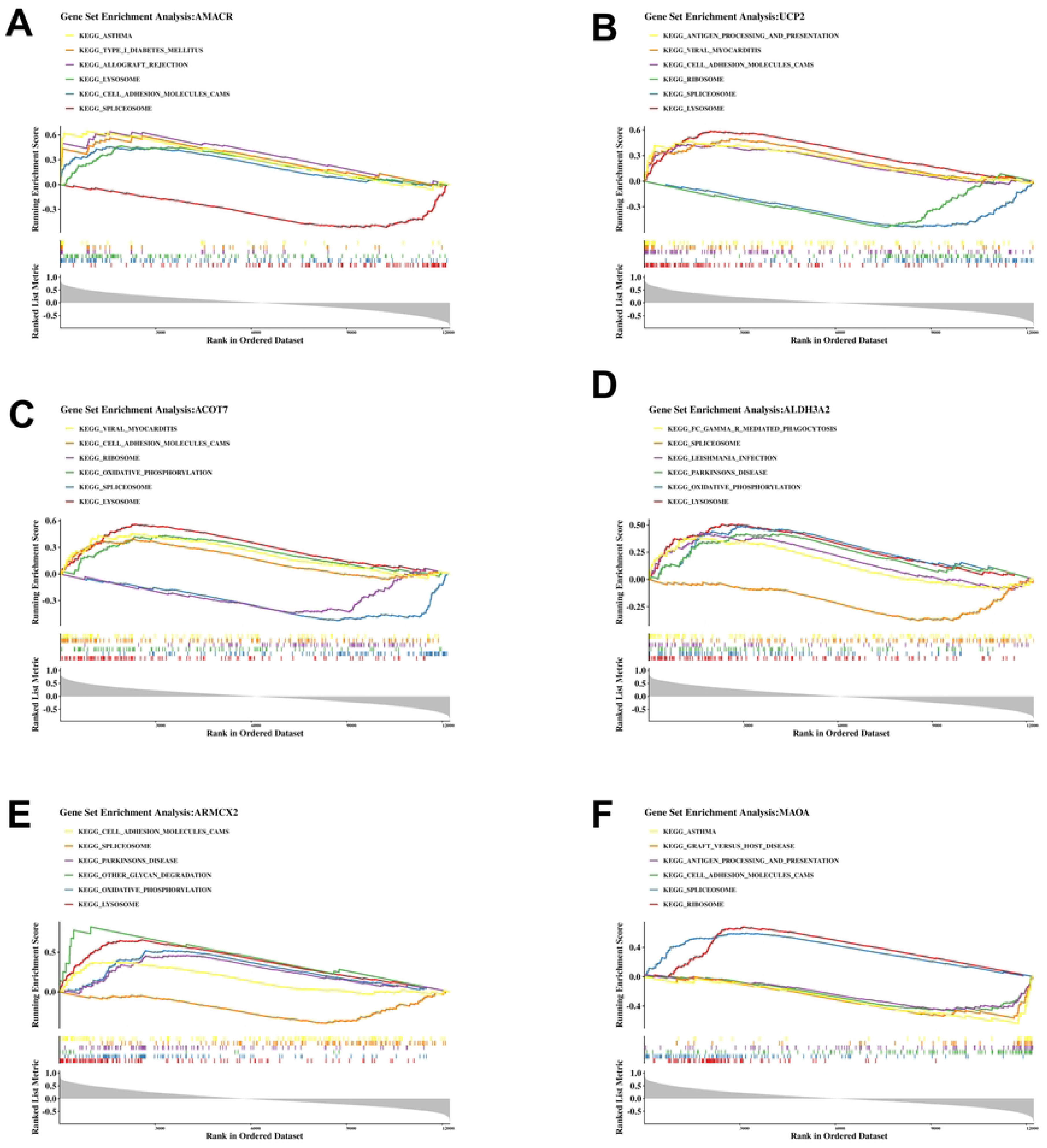
Drug-gene interaction network. Red diamonds represent hub genes. Blue circles indicate candidate drugs. Lines illustrate the predicted regulatory relationships between hub genes and candidate compounds.

### Drug Prediction and Molecular Docking Analysis

Drug-gene interaction prediction was performed based on the DSigDB database. A total of 312 non-redundant drug-gene association pairs were obtained. After drug-name standardization, 17 key candidate drugs, each targeting no fewer than three hub genes, were identified, resulting in 61 key drug–gene interaction pairs (Figure 8, Supplementary Table 5).

**Figure 8.**
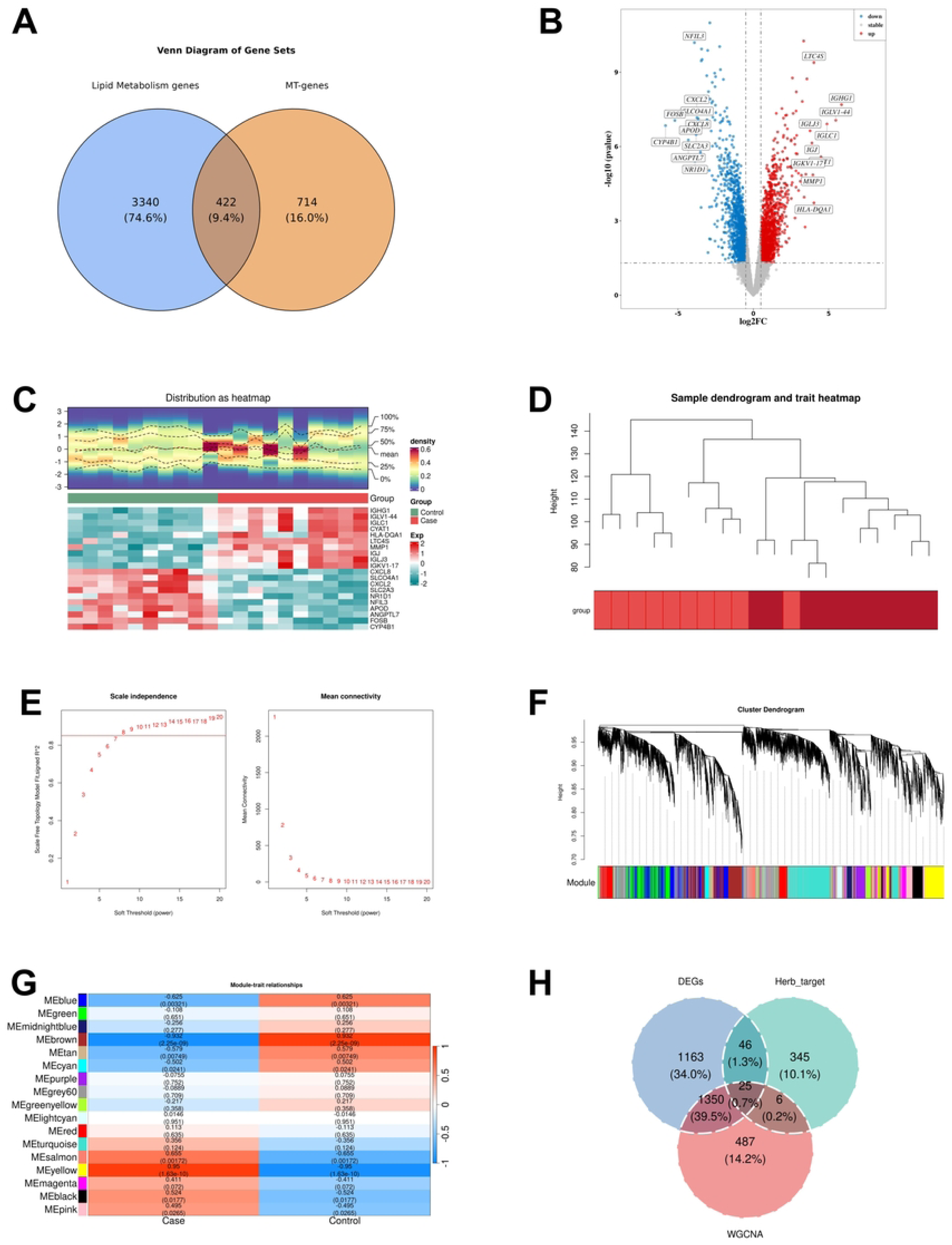
Cellular phenotypes and hub gene expression in the OA model. (A) Cell viability of chondrocytes. (B) IL-6 secretion level of chondrocytes. (C) Apoptosis analysis of chondrocytes. (D) Relative mRNA expression of six hub genes in chondrocytes. Data are presented as mean ± standard deviation. Asterisks denote statistical significance: *p < 0.05, **p < 0.01 *vs.* the Control group.

Molecular-docking simulation was performed to evaluate the binding affinity between six hub gene-encoded proteins and candidate drugs. ACOT7 exhibited the optimal binding capacity with trichostatin A, with a binding energy of −7.5 kcal/mol. ALDH3A2 showed the optimal binding capacity with trichostatin A, yielding a binding energy of −8.7 kcal/mol. AMACR possessed the optimal binding capacity with cyclosporin A at −8.3 kcal/mol. ARMCX2 obtained the optimal binding capacity with trichostatin A, with a binding energy of −6.6 kcal/mol. MAOA displayed the optimal binding capacity with quercetin, with a binding energy of −9.6 kcal/mol. UCP2 showed the optimal binding capacity with scriptaid at −8.2 kcal/mol (Supplementary Table 6, Supplementary Figure 1). The molecular-docking conformations of other candidate drugs targeting these hub genes were shown in Supplementary Figure 1.

### Cellular Phenotypes and Hub Gene Expression in the OA Model

IL-1β treatment significantly reduced chondrocyte viability compared with the Control group (Figure 8A). In the OA group, IL-6 levels were markedly elevated (Figure 8B), and the apoptotic rate of chondrocytes was also significantly increased relative to the control group (Figure 8C). Moreover, the mRNA expression levels of AMACR, UCP2, ACOT7, ALDH3A2, and ARMCX2 were significantly upregulated in the OA group compared with the Control group, whereas MAOA expression was significantly downregulated (Figure 8D).

## Discussion

By integrating omics data with gene sets related to mitochondrial and lipid metabolism, this study successfully identified six mitochondrial lipid metabolism hub genes associated with OA that exhibited diagnostic potential. These findings provide deeper insights into the mechanisms underlying mitochondrial lipid metabolism in OA and highlight clinically relevant biomarkers for the disease.

Through machine learning approaches, AMACR, UCP2, ACOT7, ALDH3A2, ARMCX2, and MAOA were identified as hub genes. Among them, the expression levels of AMACR, UCP2, ACOT7, ALDH3A2, and ARMCX2 were significantly upregulated in the OA group, whereas MAOA was significantly downregulated. Among all lipid metabolism pathways, AMACR plays a pivotal role in the β-oxidation of branched-chain fatty acids (BCFAs) within both peroxisomes and mitochondria, specifically by facilitating the stereochemical conversion of 2-methyl fatty acids^[12]^. Observations of incomplete fatty acid β-oxidation in osteoarthritic cells suggest an impairment in BCFA degradation. Consequently, we hypothesize that AMACR contributes to the initiation and progression of osteoarthritis by mediating disturbances in BCFA β-oxidation.

UCP2 is an uncoupling protein located on the inner mitochondrial membrane that counteracts oxidative stress^[13]^, which is a driving factor in OA progression. Mitochondrial dysfunction and oxidative stress are important intervention targets for age-related and degenerative diseases. By enhancing the cellular antioxidant defense system or reducing the production of mitochondrial reactive oxygen species (ROS), we can prevent the damage caused by excessive ROS, thereby maintaining normal physiological processes and tissue regeneration^[14]^.Studies have shown that palmitic acid induces chondrocyte death and the production of inflammatory mediators in articular cartilage. ACOT7 converts acyl-CoA into free fatty acids, thereby inducing apoptotic cell death in human articular chondrocytes. Elevated levels of free fatty acids (FFAs) in synovial fluid can initiate and/or exacerbate joint inflammation, potentially contributing to the onset and progression of osteoarthritis (OA)^[15]^. ALDH3A2 functions in multiple lipid metabolism pathways, catalyzing the oxidation of medium- and long-chain aliphatic aldehydes to their corresponding fatty acids. Lipid peroxidation-derived aldehyde products, such as 4-HNE, play important roles in cartilage degeneration and subchondral bone remodeling during OA progression^[16]^. The ARMCX2 protein is localized to mitochondria and is involved in the regulation of mitochondrial transport, size, and fission in neuronal axons^[17]^. To date, no systematic studies have demonstrated its specific role in OA. MAOA catalyzes the oxidative deamination of monoamine neurotransmitters. Multiple studies have shown that the degree of MAOA downregulation is negatively correlated with OA severity. Restoring MAOA expression protects against chondrocyte loss and extracellular matrix degradation through autophagy regulation[18].Taken together, these hub genes represent potential molecular markers for OA diagnosis and may participate in the progression of OA through diverse biological pathways. Furthermore, they provide a theoretical direction for the development of therapeutic strategies targeting lipid metabolism and mitochondrial function in OA.

Trichostatin A, Cyclosporin A, Quercetin, Scriptaid, and other candidate drugs exhibited low binding free energies with the proteins encoded by the hub genes. This indicates that these candidate drugs exhibit favorable binding affinities and potential regulatory interactions with the hub genes. HDACs are involved in the development of cartilage and chondrocytes and play a critical role in the pathogenesis of OA^[19]^. Previous studies have demonstrated that HDAC inhibitors can protect cartilage from disease-related damage^[19]^. Notably, the candidate drug identified in this study, Trichostatin A, is a histone deacetylase inhibitor. Therefore, Trichostatin A represents a promising therapeutic candidate for OA, exerting chondroprotective effects by targeting hub genes. Quercetin is a natural flavonoid compound with a broad range of pharmacological activities^[20]^, and it exerts significant chondroprotective effects in OA. Research has demonstrated that quercetin (QCT) significantly attenuated articular cartilage damage in osteoarthritic (OA) mice. Furthermore, QCT at a concentration of 100 µM significantly promoted the proliferation of IL-1β-stimulated chondrocytes and reduced their apoptosis^[21]^. The expression of calcineurin is elevated in OA chondrocytes. Cyclosporin A, a calcineurin inhibitor, can reduce the extent of cartilage damage. In addition, studies have shown that Cyclosporin A has mitochondrial protective functions^[22]^. Scriptaid is also a histone deacetylase (HDAC) inhibitor^[23]^. Furthermore, it has been reported that Scriptaid can protect chondrocytes by inhibiting the expression of ZNF440^[24]^. Trichostatin A, Cyclosporin A, Quercetin, and Scriptaid have been reported to exert chondroprotective effects. Furthermore, the molecular docking results in the present study indicate that these candidate drugs exhibit favorable binding affinities with OA-related hub genes. Accordingly, these compounds provide a theoretical basis for drug development targeting OA-related lipid metabolism and mitochondrial function, and hold promise as potential therapeutic candidates for OA.In this study, the bioinformatics analysis was based on public databases with a relatively limited sample size; drug prediction and molecular docking were performed as bioinformatics simulation analyses, and drug intervention experiments at the cellular or animal level have not yet been conducted. Therefore, the actual effects of the candidate drugs remain to be validated. In this study, six mitochondrial–lipid metabolism-related hub genes were identified. These genes are dysregulated in OA and demonstrate diagnostic potential for OA. These findings provide new directions for the diagnosis of OA and offer a theoretical basis for investigating the pathogenesis of OA. Subsequent studies will conduct gene intervention experiments to further elucidate the functions of each hub gene in chondrocytes, and will use cellular or animal models to further validate the hub genes and candidate drugs.

## Acknowledgments

The authors would like to thank all contributors and reviewers for their valuable comments and suggestions.

---

**Supplementary Figure 1.**
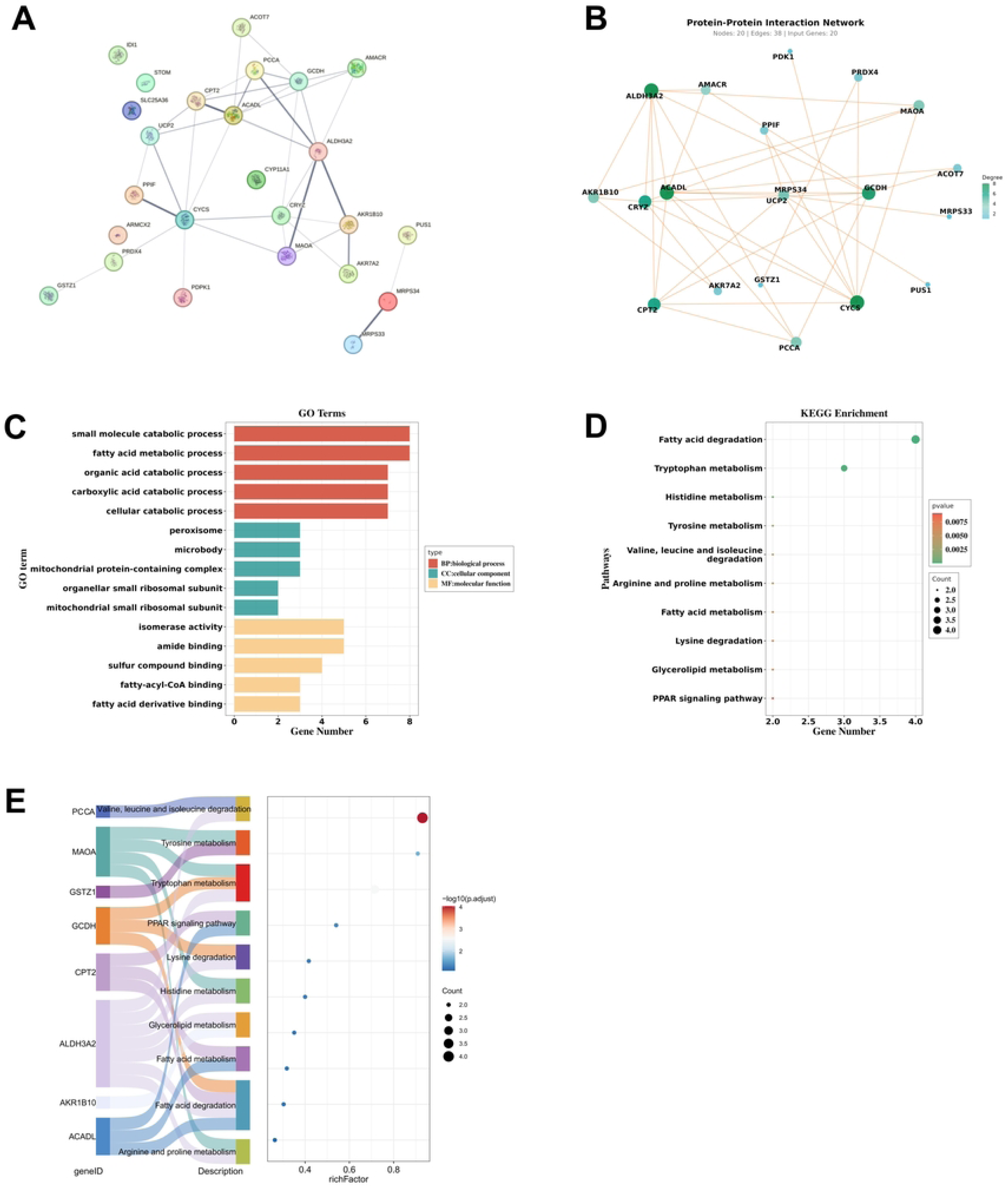
Molecular docking of candidate drugs targeting hub-gene-encoded proteins. (A) Trichostatin A (TSA)-ACOT7 (−7.5 kcal/mol). (B) Vorinostat (SAHA)-ACOT7 (−6.9 kcal/mol). (C) Valproic acid (VPA)-ACOT7 (−5.6 kcal/mol). (D) TSA-ALDH3A2 (−8.7 kcal/mol). (E) Latamoxef-ALDH3A2 (−8.5 kcal/mol). (F) Quercetin-ALDH3A2 (−7.7 kcal/mol). (G) Cyclosporin A (CsA)-AMACR (−8.3 kcal/mol). (H) Bicalutamide-AMACR (−7.6 kcal/mol). (I) Trichostatin A-AMACR (−7.0 kcal/mol). (J) Trichostatin A-ARMCX2 (−6.6 kcal/mol). (K) SAHA-ARMCX2 (−6.6 kcal/mol). (L) VPA-ARMCX2 (−4.3 kcal/mol). (M) Quercetin-MAOA (−9.6 kcal/mol). (N) TSA-MAOA (−9.4 kcal/mol). (O) VPA-MAOA (−5.8 kcal/mol). (P) Scriptaid-UCP2 (−8.2 kcal/mol). (Q) Quercetin-UCP2 (−7.6 kcal/mol). (R) VPA-UCP2 (−4.5 kcal/mol). Each subpanel shows the full protein-ligand complex and magnified local binding conformation, and indicates the binding free energy (kcal/mol).

**Supplementary Table 1:**
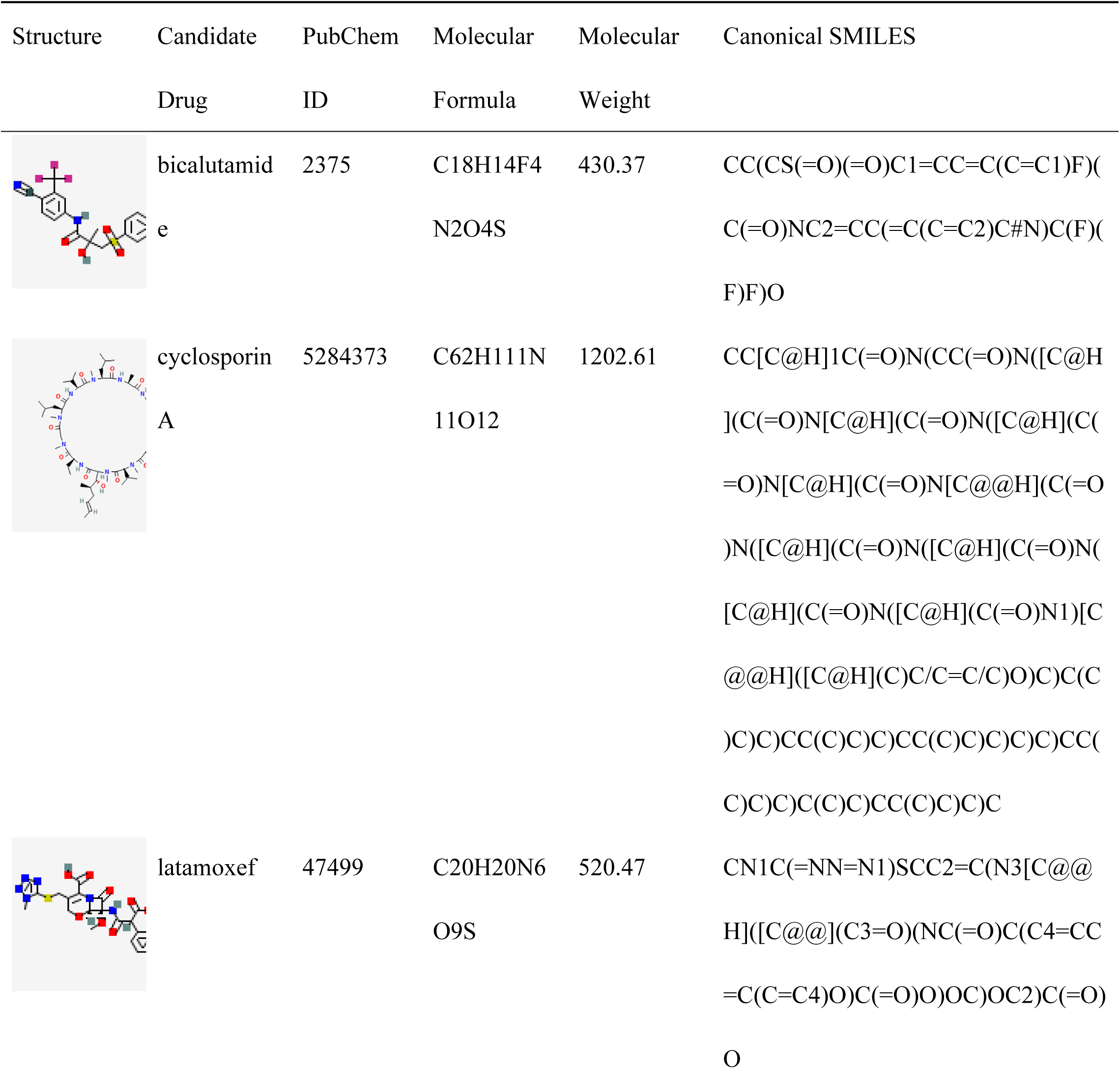
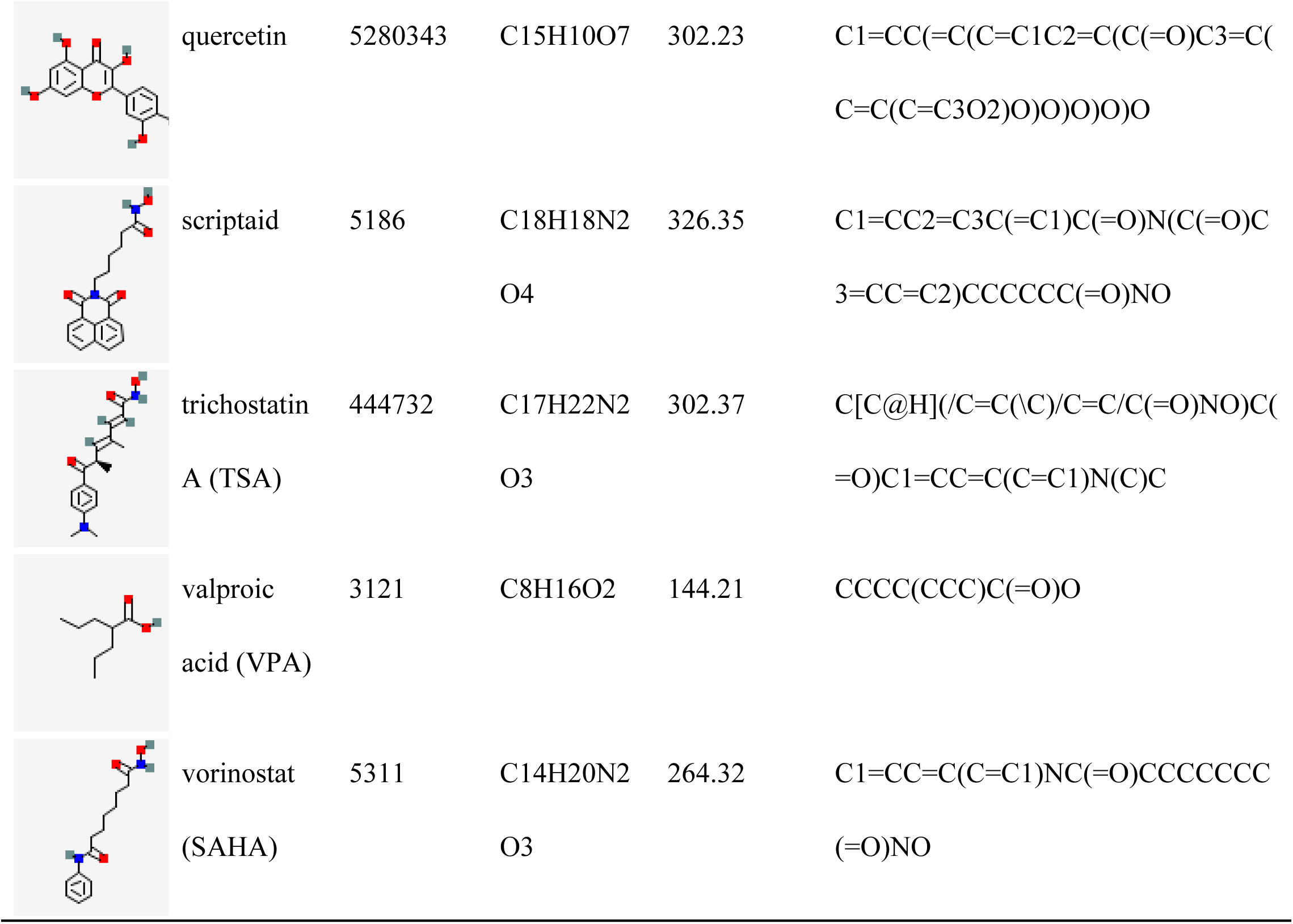
Candidate Drug Information and Their Corresponding 2D Structures.

**Supplementary Table 2:**
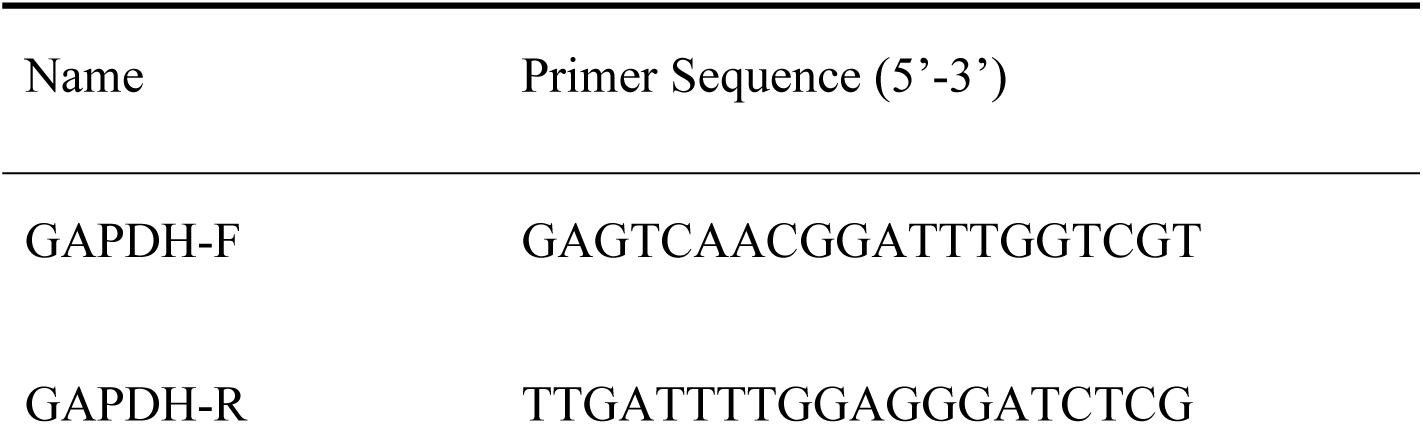
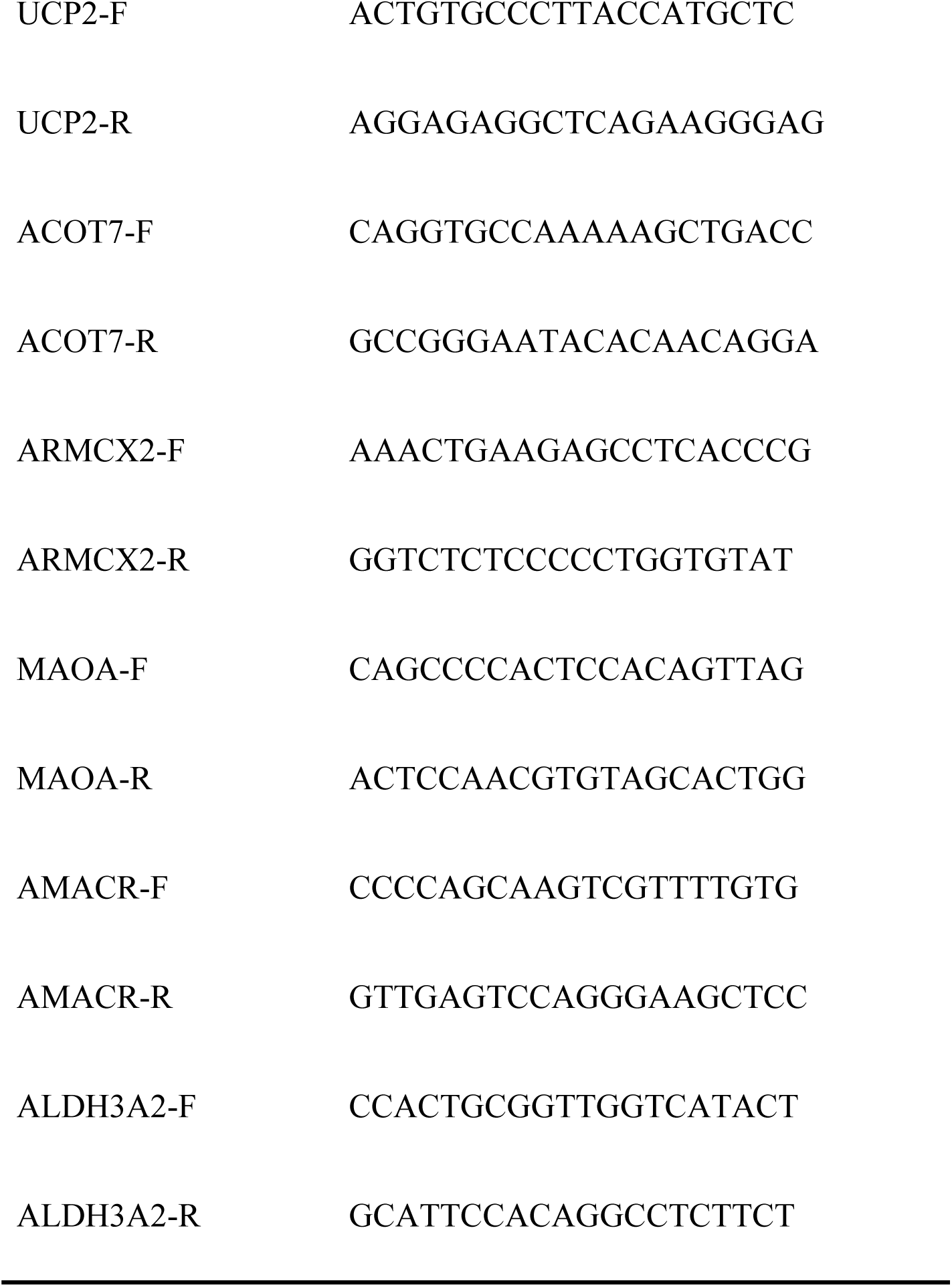
Primer Sequences for Reverse Transcription Quantitative Polymerase Chain Reaction (RT-qPCR)

**Supplementary Table 3:**
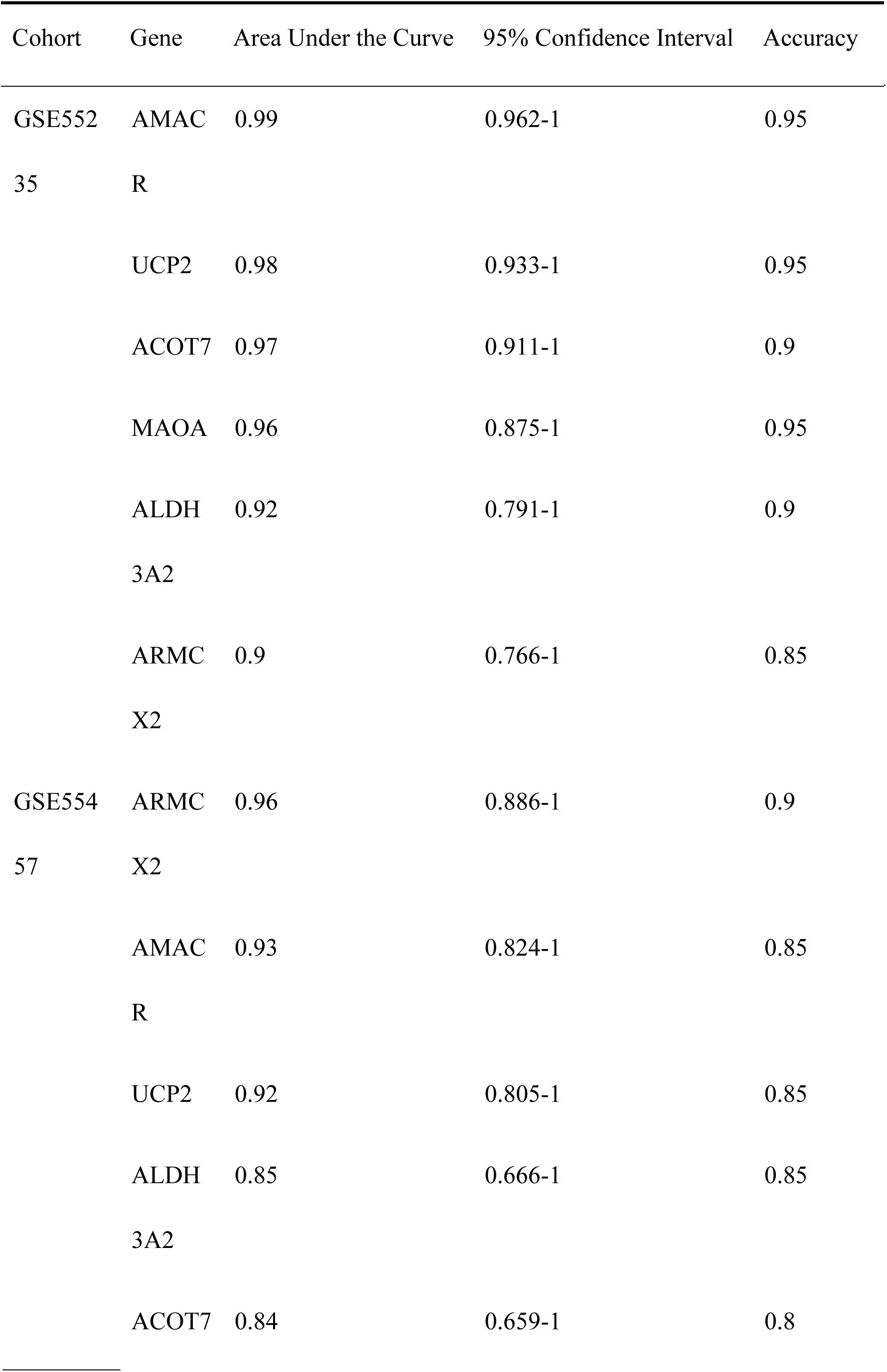
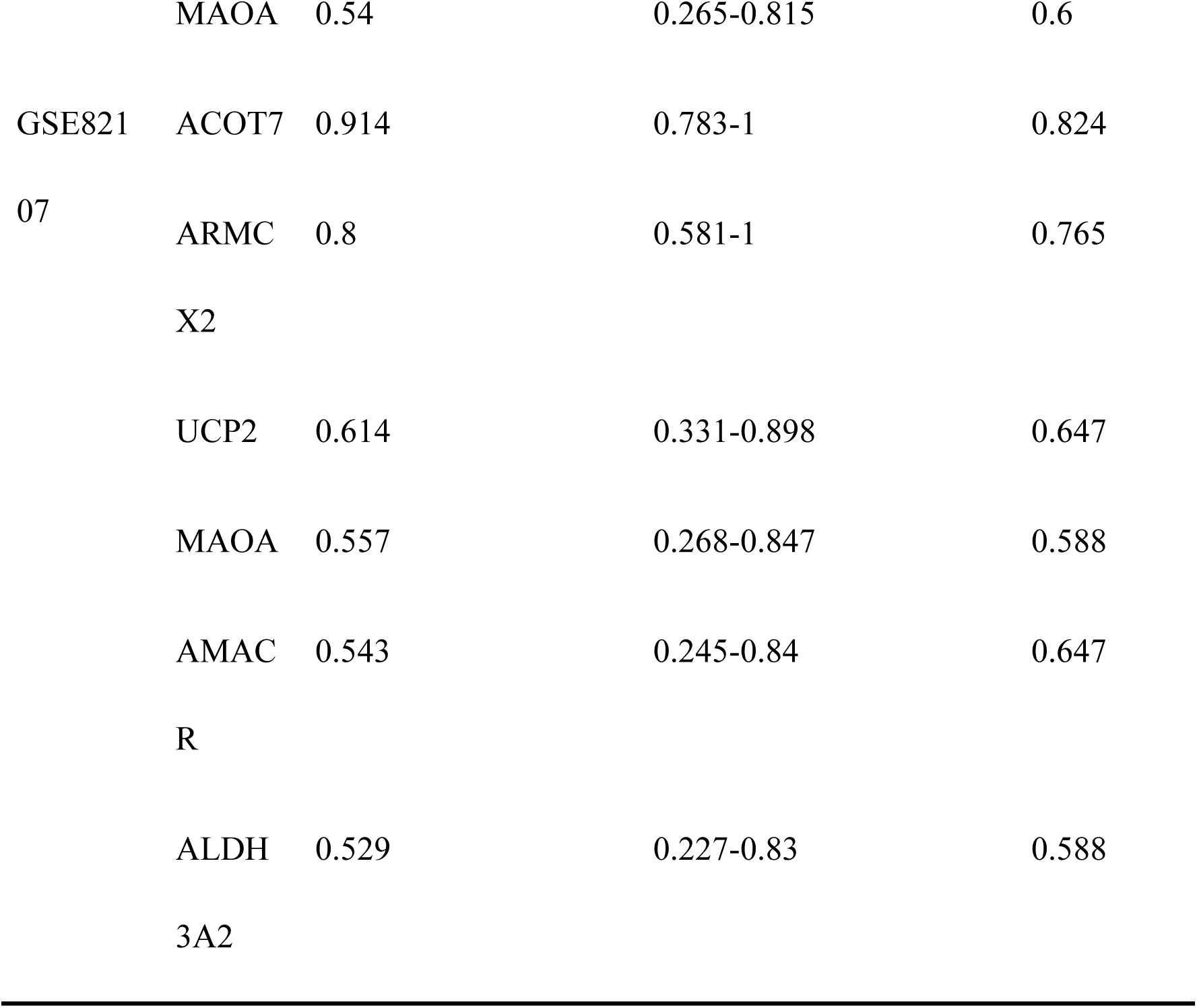
Receiver Operating Characteristic (ROC) Analysis Results of Hub Genes.

**Supplementary Table 4:** Top 10 Kyoto Encyclopedia of Genes and Genomes Pathway Enrichment Results.

| GeneSet | Pathway Short | Normalized | Adjusted | setSize | Direction |
| --- | --- | --- | --- | --- | --- |
|  |  | Enrichment Score | p-value |  |  |
| ACOT7 | LYSOSOME | 2.596 | 1.83e-08 | 108 | Positive |
| ACOT7 | SPLICEOSOME | -2.410 | 5.45e-08 | 104 | Negative |
| ACOT7 | OXIDATIVE PHOSPHORYLATION | 1.952 | 1.05e-03 | 87 | Positive |
| ACOT7 | RIBOSOME | -1.934 | 1.05e-03 | 82 | Negative |
| ACOT7 | CELL ADHESION MOLECULES CAMS | 1.833 | 1.05e-03 | 118 | Positive |
| ACOT7 | VIRAL MYOCARDITIS | 1.938 | 2.23e-03 | 66 | Positive |
| ACOT7 | ECM RECEPTOR INTERACTION | 1.824 | 5.84e-03 | 81 | Positive |
| ACOT7 | OTHER GLYCAN DEGRADATION | 1.967 | 1.40e-02 | 13 | Positive |
| ACOT7 | ASTHMA | 1.885 | 1.59e-02 | 27 | Positive |
| ACOT7 | SYSTEMIC LUPUS ERYTHEMATOSUS | 1.751 | 2.17e-02 | 52 | Positive |
| ALDH3A2 | LYSOSOME | 2.432 | 2.30e-08 | 108 | Positive |
| ALDH3A2 | OXIDATIVE PHOSPHORYLATION | 2.248 | 1.03e-05 | 87 | Positive |
| ALDH3A2 | PARKINSONS DISEASE | 1.924 | 3.48e-03 | 87 | Positive |
| ALDH3A2 | LEISHMANIA INFECTION | 1.919 | 8.59e-03 | 69 | Positive |
| ALDH3A2 | SPLICEOSOME | -1.779 | 8.82e-03 | 104 | Negative |
| ALDH3A2 | FC GAMMA R MEDIATED<br>PHAGOCYTOSIS | 1.802 | 1.00e-02 | 88 | Positive |
| ALDH3A2 | PRIMARY IMMUNODEFICIENCY | 1.879 | 3.45e-02 | 34 | Positive |
| ALDH3A2 | ARACHIDONIC ACID METABOLISM | 1.827 | 3.45e-02 | 48 | Positive |
| ALDH3A2 | BLADDER CANCER | -1.791 | 3.45e-02 | 41 | Negative |
| ALDH3A2 | PEROXISOME | 1.664 | 3.45e-02 | 70 | Positive |
| AMACR | SPLICEOSOME | -2.353 | 1.29e-07 | 104 | Negative |
| AMACR | CELL ADHESION MOLECULES CAMS | 2.207 | 8.58e-07 | 118 | Positive |
| AMACR | LYSOSOME | 2.226 | 2.32e-06 | 108 | Positive |
| AMACR | ALLOGRAFT REJECTION | 2.374 | 2.69e-05 | 34 | Positive |
| AMACR | TYPE I DIABETES MELLITUS | 2.267 | 2.69e-05 | 40 | Positive |
| AMACR | ASTHMA | 2.265 | 2.27e-04 | 27 | Positive |
| AMACR | INTESTINAL IMMUNE NETWORK<br>FOR IGA PRODUCTION | 2.099 | 5.71e-04 | 42 | Positive |
| AMACR | LEISHMANIA INFECTION | 2.036 | 6.57e-04 | 69 | Positive |
| AMACR | SYSTEMIC LUPUS ERYTHEMATOSUS | 2.033 | 9.43e-04 | 52 | Positive |
| AMACR | GRAFT VERSUS HOST DISEASE | 2.065 | 1.67e-03 | 37 | Positive |
| ARMCX2 | LYSOSOME | 2.999 | 1.83e-08 | 108 | Positive |
| ARMCX2 | OXIDATIVE PHOSPHORYLATION | 2.302 | 1.32e-06 | 87 | Positive |
| ARMCX2 | OTHER GLYCAN DEGRADATION | 2.300 | 1.18e-04 | 13 | Positive |
| ARMCX2 | PARKINSONS DISEASE | 2.028 | 2.95e-04 | 87 | Positive |
| ARMCX2 | SPLICEOSOME | -1.961 | 6.42e-04 | 104 | Negative |
| ARMCX2 | CELL ADHESION MOLECULES CAMS | 1.753 | 2.76e-03 | 118 | Positive |
| ARMCX2 | ALZHEIMERS DISEASE | 1.790 | 3.48e-03 | 132 | Positive |
| ARMCX2 | GLYCOSAMINOGLYCAN<br>DEGRADATION | 2.027 | 6.05e-03 | 21 | Positive |
| ARMCX2 | VIRAL MYOCARDITIS | 1.873 | 6.05e-03 | 66 | Positive |
| ARMCX2 | FC GAMMA R MEDIATED<br>PHAGOCYTOSIS | 1.757 | 7.89e-03 | 88 | Positive |
| MAOA | RIBOSOME | 2.867 | 9.15e-09 | 82 | Positive |
| MAOA | SPLICEOSOME | 2.612 | 9.15e-09 | 104 | Positive |
| MAOA | CELL ADHESION MOLECULES CAMS | -2.225 | 6.77e-07 | 118 | Negative |
| MAOA | ANTIGEN PROCESSING AND<br>PRESENTATION | -2.057 | 2.92e-04 | 79 | Negative |
| MAOA | GRAFT VERSUS HOST DISEASE | -2.122 | 5.81e-04 | 37 | Negative |
| MAOA | ASTHMA | -2.215 | 6.65e-04 | 27 | Negative |
| MAOA | ECM RECEPTOR INTERACTION | -1.967 | 6.65e-04 | 81 | Negative |
| MAOA | INTESTINAL IMMUNE NETWORK<br>FOR IGA PRODUCTION | -2.094 | 6.74e-04 | 42 | Negative |
| MAOA | SYSTEMIC LUPUS ERYTHEMATOSUS | -2.096 | 6.82e-04 | 52 | Negative |
| MAOA | ALLOGRAFT REJECTION | -2.184 | 9.01e-04 | 34 | Negative |
| UCP2 | LYSOSOME | 2.655 | 9.15e-09 | 108 | Positive |
| UCP2 | SPLICEOSOME | -2.500 | 9.15e-09 | 104 | Negative |
| UCP2 | RIBOSOME | -2.429 | 1.77e-07 | 82 | Negative |
| UCP2 | CELL ADHESION MOLECULES CAMS | 1.983 | 4.70e-05 | 118 | Positive |
| UCP2 | VIRAL MYOCARDITIS | 2.048 | 2.50e-04 | 66 | Positive |
| UCP2 | ANTIGEN PROCESSING AND<br>PRESENTATION | 2.004 | 5.56e-04 | 79 | Positive |
| UCP2 | ASTHMA | 2.157 | 1.14e-03 | 27 | Positive |
| UCP2 | LEISHMANIA INFECTION | 1.918 | 1.14e-03 | 69 | Positive |
| UCP2 | SYSTEMIC LUPUS ERYTHEMATOSUS | 1.930 | 2.33e-03 | 52 | Positive |
| UCP2 | ALLOGRAFT REJECTION | 2.027 | 2.92e-03 | 34 | Positive |

**Supplementary Table 5:**
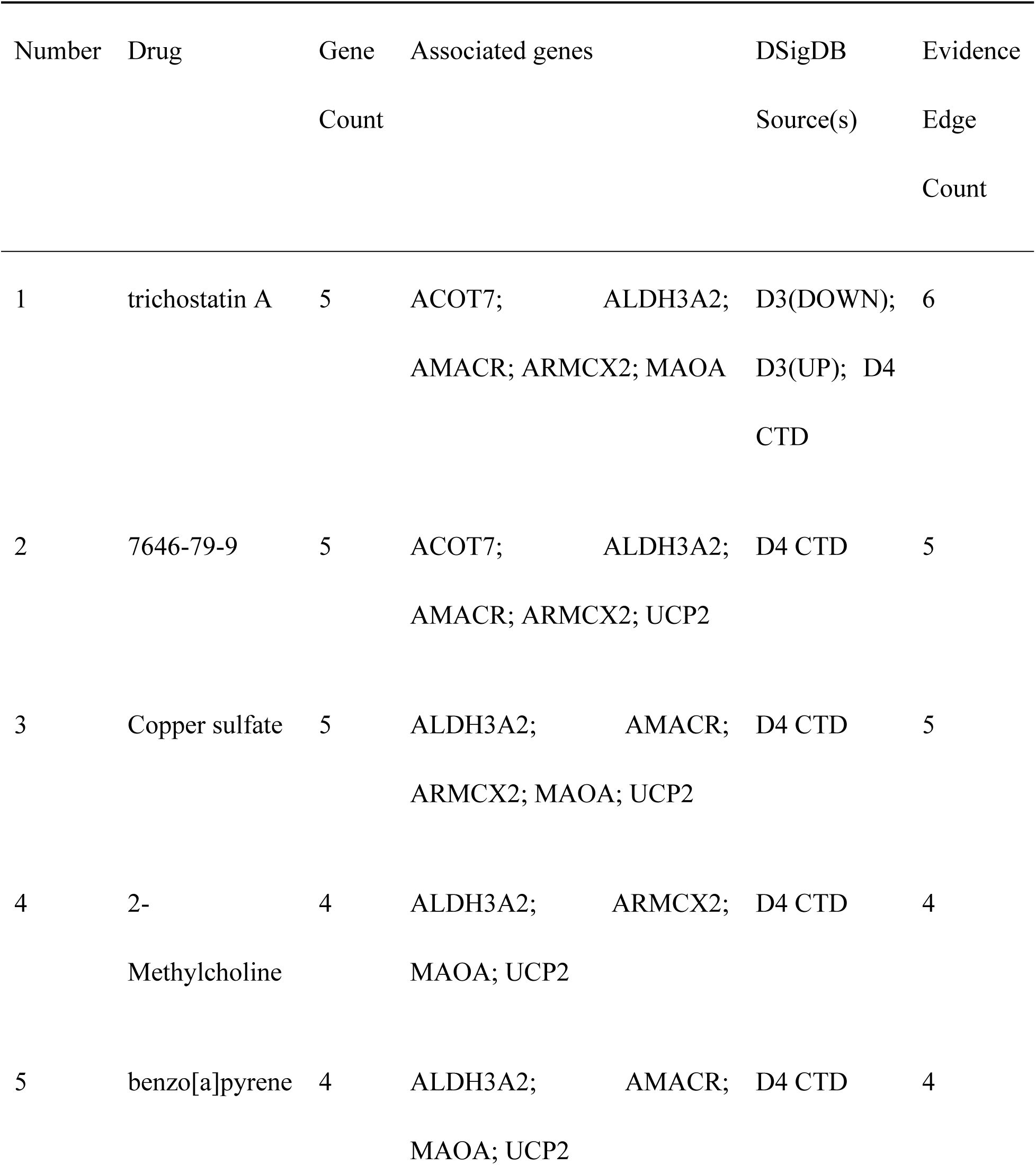
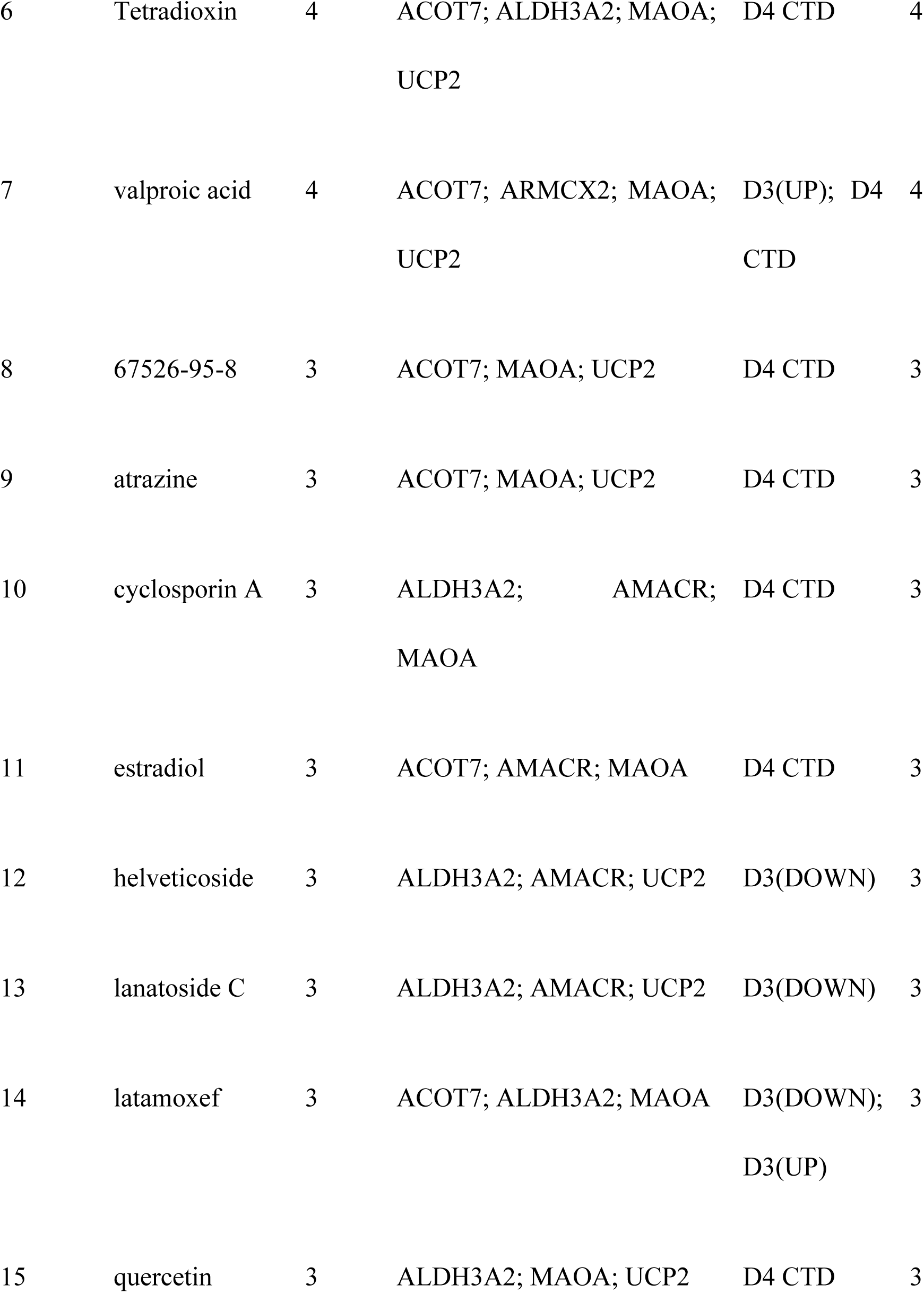
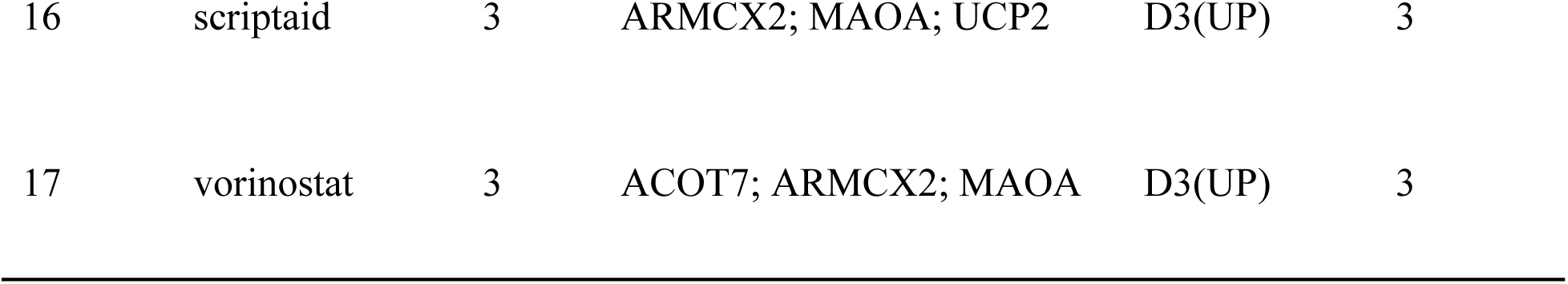
Candidate Drugs Targeting Hub Genes.

**Supplementary Table 6:**
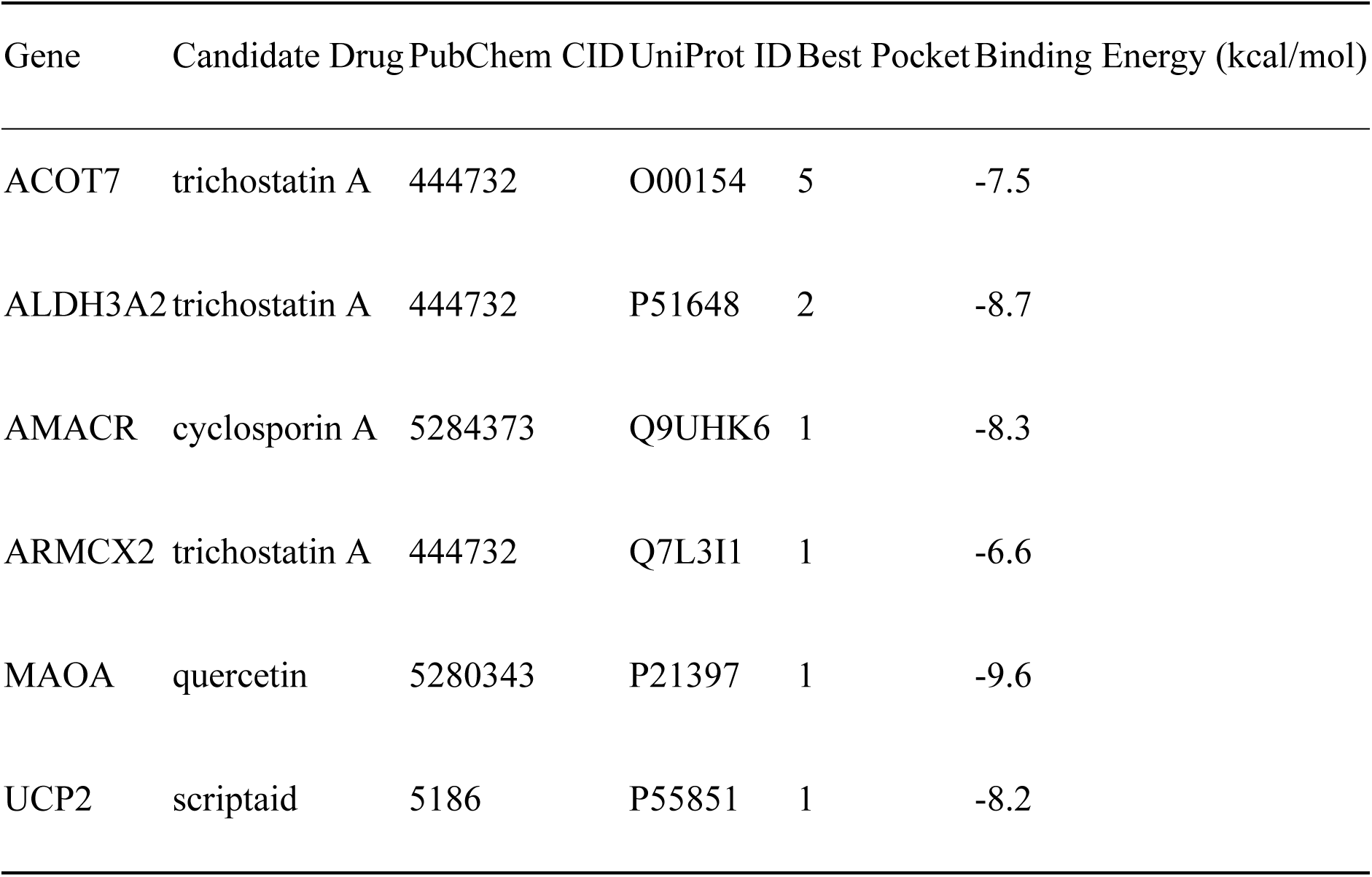
Information on the Drugs and Proteins Encoded by Hub Genes.

